# Identifying endogenous substrates of the 26S proteasome through site-specific photocrosslinking

**DOI:** 10.64898/2026.07.19.739412

**Authors:** Santiago Yori Restrepo, Andreas Martin

## Abstract

The 26S proteasome is the hub for regulated protein turnover in eukaryotic cells. Degradation of proteins by the Ubiquitin-Proteasome System plays critical roles in every aspect of cell biology, such as the regulation of gene transcription, the quality control of translation and protein folding, and protein transport across membranes. While mRNA levels and protein abundances can be readily measured with a robust set of established tools, only a few methodologies exist to identify proteins that are degraded by the proteasome rather than the lysosome as the second major pathway for turnover. Here, we sought to address this by using genetic code expansion to introduce a photo-crosslinkable unnatural amino acid into the yeast 26S proteasome and capture cellular protein substrates as they translocate through the proteasomal ATPase motor. *In vitro* biochemical experiments confirmed that these modified proteasomes are functional, which allowed us to introduce them into live yeast cells for *in vivo* crosslinking and the identification of enriched ATP-dependent substrates by mass spectrometry. These experiments revealed a very diverse pool of proteasomal substrates that markedly changed upon cell exposure to endoplasmic reticulum stress. Together, our results represent an exciting avenue for probing the landscape of proteasomal substrates and its changes in response to various cellular conditions and stresses.

## Introduction

An estimated 80% of eukaryotic proteins meet their end through degradation by the 26S proteasome as the final stage within the Ubiquitin-Proteasome System (UPS) ^1^. This fact highlights an incredible feature of this essential proteolytic complex: its activity must be both promiscuous and highly selective, as it degrades the bulk of the cellular proteome, while also picking out regulatory proteins with single digit copy numbers among millions of candidates ^2^. Key in this balance are the specific protein recruitment to the proteasome and the engagement by the proteasomal ATPase motor. For the majority of substrates this is accomplished by a bipartite degradation signal that consists of an enzymatically attached ubiquitin chain for targeting and an intrinsic unstructured initiation region for insertion into the proteasome’s central channel ^3^. Hundreds of E3 ubiquitin ligases, together with dozens of E2 ubiquitin conjugating enzymes, dictate the specificity of substrate selection as well as the length and type of ubiquitin chains that are usually attached to lysine residues of the substrate ^4^, but there is also a growing number of identified ubiquitin-independent targeting mechanisms ^5–9^.

Flexible initiation regions that can be engaged by the AAA+ (ATPases associated with various cellular activities) ATPase of the proteasome may be intrinsic to the substrate structure or be generated through the action of another AAA+ motor, in many cases the cytosolic protein unfoldase CDC48 (p97/VCP in humans) acting upstream of the 26S proteasome ^10–12^.

Selectivity in degradation is accomplished by the proteasome’s architecture as a compartmental protease, with the proteolytic active sites sequestered in a gated internal chamber of the 20S core peptidase (CP) that is composed of 14 distinct subunits, α1-7 and β1-7 (Fig. 1A, gray) ^13^. Although there is evidence for unstructured polypeptides diffusing directly into isolated 20S CP ^14^, protein degradation by the proteasome is mostly regulated by ATP-dependent or -independent “caps” that bind to one or both sides of the 20S CP to control the apical gates for substrate access ^15^. The most prominent of these caps is the 19S regulatory particle (RP), which uses ATP hydrolysis to mechanically unfold and translocate substrates into the 20S CP degradation chamber. This 19S RP can be biochemically divided into two subcomplexes, the base and the lid ^16^. The base contains the large scaffolding subunit Rpn2, six distinct ATPase subunits, Rpt1-Rpt6, that form the heterohexameric AAA+ motor of the proteasome ^17,18^, and three ubiquitin receptors, Rpn1, Rpn10, and Rpn13, for ubiquitin-dependent substrate recruitment (Fig. 1A) ^19–21^. Each Rpt subunit consists of an N-terminal domain and a C-terminal AAA+ ATPase domain, which in the hexamer form two stacked rings. While the N-terminal domain ring (N-ring) is rigid and static, the AAA+ ring uses the energy of ATP hydrolysis to undergo conformational changes that generate the mechanical force for substrate unfolding and translocation through the central channel into the 20S CP. Conserved pore-1 and pore-2 loops that protrude from each Rpt subunit into the channel thereby mediate steric contacts to the substrate polypeptide, with the pore-1 loop’s highly conserved Tyr or Phe providing the main grip for translocation ^22^. The lid subcomplex consists of 9 subunits, which include the essential Zn^2+^-dependent deubiquitinase (DUB) Rpn11 that removes ubiquitin modifications from substrates in a co-translocational manner (Fig. 1A, orange) ^23–26^.

**Figure 1.**
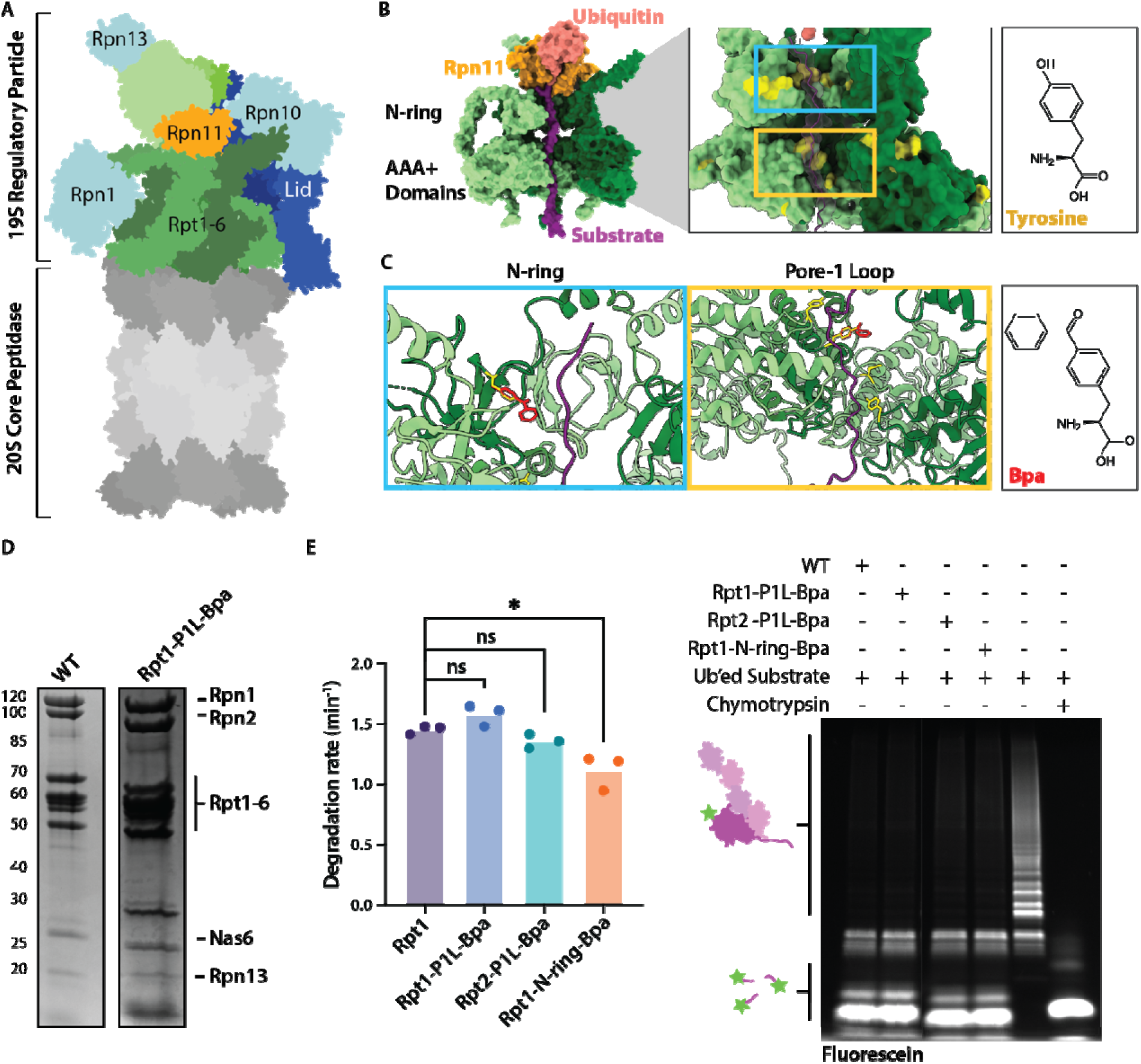
Structure-guided introduction of UV-activated crosslinkable Bpa. **A.** Architecture of the yeast 26S proteasome (PDB ID: 4CR4). The 20S core peptidase is shown in gray, capped by the colored 19S regulatory particle. The ATPase subunits Rpt1-Rpt6 of the base subcomplex are shown in alternating medium and dark green, the ubiquitin-receptor subunits Rpn1, Rpn10, and Rpn13 are shown in cyan, and the large scaffolding subunit Rpn2 in light green. The Rpn11 deubiquitinase subunit of the lid subcomplex is depicted in orange and other lid subunits in medium and dark blue. **B.** Left: Structure of the ATPase motor (green) with an engaged substrate polypeptide (purple) in the central channel and the first, substrate-attached ubiquitin (salmon) bound to Rpn11 (orange). Two regions in the Rpt hexamer, the N-ring and the AAA+ domains, are in close contact with the substrate polypeptide. Right: Surface representation of the Rpt1-Rpt6 hexamer with tyrosine residues highlighted in yellow. The central channel through the N-ring and the AAA+ motor domains is highlighted with a cyan and yellow box, respectively. **C.** Zoom-in on the central channel through the N-ring (left) and AAA+ motor domains (right), with tyrosine residues shown in yellow stick representations and models for Tyr-to-Bpa substitutions shown in red. **D.** Representative Coomassie-stained SDS-PAGE gel showing the purified wild-type and Bpa-containing base subcomplexes recombinantly expressed in *E. coli*. **E.** Left: Rates for the multiple-turnover degradation of a ubiquitinated titin model substrates by 26S proteasomes reconstituted with the wild-type base subcomplex or Bpa-containing variants. Shown are the averages for n = 3 technical replicates. Statistical significance was calculated using a one-way ANOVA test: * p<0.05; ns, p>0.05. Right: In-gel fluorescence detection of the fluorescein-labelled, ubiquitinated substrate and peptide products present at the endpoints of degradation reactions with wild-type and Rpt1-PL1-Bpa or Rpt2-PL1-Bpa containing proteasomes.

Understanding the vast pool of cellular substrates that are degraded by the 26S proteasome has been a long-standing goal of the UPS research community. However, previous attempts in elucidating the proteasomal “degradome” were hampered by a considerable crosstalk between proteasomal and lysosomal turnover, and numerous feedback mechanisms that are activated in response to inhibiting either one of these principal degradation pathways. Earlier important work aimed at understanding the ubiquitinated proteome ^27^, while more recent studies used crosslinking to the 20S CP in cell lysates for identifying potential proteasome substrates in bulk ^28^. However, we were still lacking a specific method to survey the hundreds or thousands of substrates that are specifically degraded by the 26S proteasome in a 19S RP-mediated, ATP-hydrolysis-dependent manner.

We therefore sought to develop an approach in which committed substrates are trapped within the AAA+ ATPase motor as they pass through the central channel of 26S proteasomes in living cells. Here, we demonstrate that genetic code expansion, incorporation of the unnatural amino acid p-benzoyl-L-phenylalanine (Bpa), and photo-induced crosslinking can serve as a potent strategy to capture substrates on their way to proteasomal degradation for mass-spectrometric identification. We used biochemical reconstitution experiments to confirm that the structure-guided incorporation of Bpa in the AAA+ motor results in crosslinkable, yet functional proteasomes that can be implemented in living cells. Furthermore, we show that perturbing protein folding in the endoplasmic reticulum causes major changes in the proteasome substrate pool, indicating the large potential of our crosslinking approach for studying shifts in proteasomal degradation in response to cellular stresses or needs.

## Results

### Structure-guided introduction of Bpa into the proteasomal motor

Prior work leveraged our recombinant expression systems for the lid and base subcomplexes of the yeast 19S RP to incorporate the unnatural amino acid (UAA) p-azido-L-phenylalanine (AzF) by amber stop codon suppression, attach fluorescent dyes, and monitor substrate degradation as well as conformational changes through fluorescence and FRET ^29–31^. For crosslinking of committed substrates during degradation, we turned our attention to the UAA p-benzoyl-L-phenylalanine (Bpa) due to its useful chemical properties, including the activation to a diradical triplet state by 365 nm light, the non-specific crosslinking to C-H bonds in nearby proteins, and the ability to reactivate Bpa after relaxation of the triplet state for repeated crosslinking attempts ^32–34^ (Supp. Fig. 1). We envisioned a strategy where we could trap substrates in the central channel of the proteasome, after they were committed to degradation, yet prior to their proteolytic cleavage within the 20S core peptidase. To choose a position for the placement of Bpa within the *S. cerevisiae* 26S proteasome’s 33 distinct subunits, we consulted the cryo-EM structures of the substrate-engaged yeast proteasome ^35^ (Fig. 1B). The structures show that the substrate on its path through the 19S RP comes within 6 Å of three positions in the ATPase hexamer: the N-ring, the pore-1 loops, and the pore-2 loops. While there was an abundance of tyrosine residues to be replaced with Bpa along the substrate path, we were particularly optimistic about the pore-1 loop Tyr, as this residue is conserved in all Rpt subunits and acting akin a gear tooth to propel the substrate (Supp. Fig. 2A). Modeling a Tyr-to-Bpa replacement into the cryo-EM structure confirmed that the central channel of the AAA+ hexamer could accommodate the slightly larger residue (Fig. 1C). Additionally, past mutational and biochemical studies with reconstituted proteasomes revealed that the Rpt subunits have unequal contributions to unfolding and degradation, with the proteasome variant containing a Tyr-to-Ala mutation in Rpt1 retaining near wild-type activity ^18^. We therefore hypothesized that introducing a Bpa residue in this position may maintain native proteasomal function. To test this, the recombinant expression system for the base subcomplex was modified to enable the incorporation of Bpa in Rpt1 and Rpt2 at various positions along the substrate path ^36^. Proper expression and assembly of the base variants was initially assessed by size-exclusion chromatography and SDS-PAGE (Fig. 1D; Supp. Fig. 2B). Of the variants tested, three appeared to assemble correctly: Bpa incorporated in the N-domain of Rpt1 (Y181Bpa, named Rpt1-N-ring-Bpa), in the pore-1 loop of Rpt1 (Y283Bpa, Rpt1-P1L-Bpa), or in the pore-1 loop of Rpt2 (Y256Bpa, Rpt2-P1L-Bpa). In contrast, mutating pore-2 loop positions in Rpt1 (A323Bpa or G325Bpa) exhibited poor to no expression as seen by SDS-PAGE (Supp. Fig. 2B).

The three purified base variants with successful Bpa incorporations were *in vitro* reconstituted with purified 19S lid and 20S CP to form crosslinking-competent 26S proteasomes. To assay the activity of these Bpa-containing proteasome variants, we measured their degradation of a model substrate, in which the titin-I27 domain containing a destabilizing V15P mutation (I27^V15P^) was fused to a C-terminal disordered tail of 35 residues derived from cyclin-B (Supp. Fig. 3A). This model substrate was *in vitro* ubiquitinated on a single Lys residue in its flexible tail for proteasome targeting and sortase-labeled at its N-terminus with fluorescein amidite (FAM) for monitoring degradation by SDS-PAGE and fluorescence polarization (Supp. Fig. 3B). Analysis by in-gel fluorescence showed that all assembled base variants could degrade the ubiquitinated model substrate (Fig. 1E). Polarization measurement revealed a degradation rate of 1.58 min^-1^ for the Rpt1-P1L-Bpa variant, similar to the wild-type activity of 1.45 min^-1^, while Rpt2-P1L-Bpa and Rpt1-N-ring-Bpa showed slightly lower rates of 1.35 min^-1^ and 1.11 min^-1^, respectively (Fig. 1E; Supp. Fig. 3B). Interestingly, those degradation rates roughly correlated with the observed assembly efficiencies of the base variants, potentially indicating the overall “fitness” of those complexes (Supp. Fig. 2B). We were surprised to see a slight defect in degradation for the Rpt1-N-ring-Bpa variant. Modeling of the Bpa at this position shows that the benzophenone side chain protrudes only marginally into the pore of the N-ring with no blockage of the substrate path (Supp. Fig. 4), but it is possible that a partial steric obstruction paired with the rigidity of the N-ring results in compromised substrate entry into the motor for engagement. We therefore decided to proceed with our pore-1 loop variants, given their similarities in degradation activities with the wild-type proteasome. Furthermore, the pore-1 loop is much further down in the central channel than the N-ring position and therefore well shielded from proteins that may bind to the proteasome surface without being committed to degradation.

### Crosslinks to 26S proteasomes are UV-dependent and specific to engageable substrates

Having confirmed that the Bpa-containing proteasomes are active in substrate degradation, we aimed to test their UV-dependent crosslinking. Reconstituted degradation reactions were allowed to proceed either in the dark or exposed to 365 nm light for 15 minutes, before reactions were separated by SDS-PAGE and analyzed by fluorescence and Western blotting against the crosslinking subunit using anti-Rpt1 or anti-Rpt2 antibodies. Initial crosslinking experiments with the Rpt1-P1L-Bpa and Rpt2-P1L-Bpa variants showed the Rpt1/2 band on the SDS-PAGE gel shifting to a single band of >120 kDa, although the proteasomes were incubated with the polyubiquitinated model substrate that itself migrates as a laddered smear on the gel (Supp. Fig. 3A). The approximate size of this UV-dependent band suggested that Rpt1/2-P1L-Bpa were forming a covalent adduct with an adjacent Rpt subunit. An unrelated experiment, aimed at assessing whether illumination by the FPLC’s UV detector could initiate crosslinking during purification, also showed a ∼120 kDa band after exposure to 365 nm light and supported our inter-Rpt crosslinking hypothesis (Supp. Fig 2C). We hypothesized that the consumption of our model substrate during the 15 min incubation for crosslinking would leave a substantial fraction of proteasomes substrate-free and thus more prone to self-crosslinking, especially when considering the proximity of neighboring pore-1 loops in the spiral-staircase arrangement of Rpt subunits (Fig. 1C) ^35^. To increase the probability for substrate-crosslinks in these proof-of-principle experiments, we therefore decided to stall translocation by inhibiting substrate de-ubiquitination with a catalytically dead deubiquitinase mutant Rpn11^AXA^ (Figure 2A) ^25,37^. As expected, incorporation of the Rpn11^AXA^ mutant in proteasome variants resulted in impaired degradation and accumulation of ubiquitinated substrates, as seen by a gel-based endpoint assay (Figure 2B, AXA lanes). Importantly, Western blotting for Rpt1 and Rpt2 revealed an additional stall-dependent shift of the Rpt bands into a higher molecular-weight smear, corresponding to UV-dependent crosslinking of the ATPase subunit to the ubiquitinated model substrate (Figure 2B, bottom right; Supp. Fig. 5).

**Figure 2.**
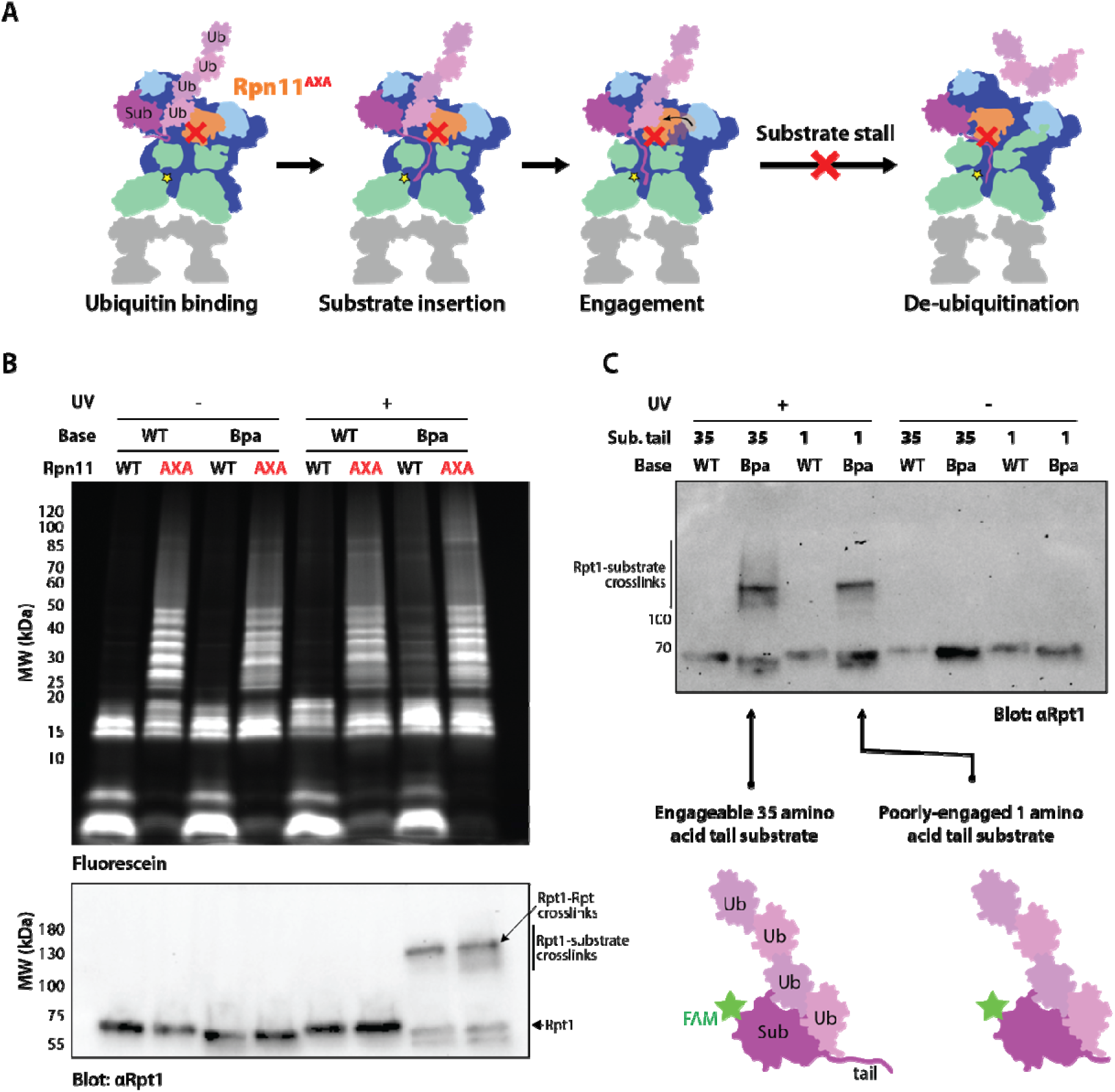
UV-dependent substrate crosslinking of Bpa-containing 26S proteasomes. **A.** Schematic of the early steps in degradation of a ubiquitinated substrate (purple) by the 26S proteasome. The AXA mutation in Rpn11 inhibits substrate deubiquitination (indicated by the red cross) and leads to a stall of substrate processing after substrate tail insertion and engagement by the ATPase motor, during which the proteasome switches from the resting to processing conformational states. **B.** Top: Monitoring of substrate in-gel fluorescence demonstrates that the Rpn11^AXA^ variant prevents degradation and the formation of peptide products (AXA lanes). Upon illumination with UV light, Bpa-containing proteasomes with wild-type Rpn11 also show some substrate stabilization due to crosslinking of Rpt1. Bottom: Blotting for Rpt1 shows a UV-dependent shift for Bpa-containing Rpt1 to higher-molecular weight. For proteasomes with the Rpn11^AXA^ mutant and consequently stalled substrate, UV irradiation results in smear of crosslinks between Rpt1 and polyubiquitinated substrate. **C.** Substrate crosslinking requires an initiation region of sufficient length. Incubation with an engagement-incompetent 1-mer tailed substrate prevents the formation of a ubiquitin-substrate smear for Rpt1 and only leads to crosslink between Rpt1 and another Rpt.

Because the proteasome functions as a hub for many transiently interacting cofactors, adaptors, shuttle receptors, and other proteins, we wanted to ensure that our Bpa placement allows selective crosslinking only to committed substrates that are translocating through the ATPase motor. We therefore tested crosslinking to an engagement-incompetent substrate variant that contained only a single-amino acid tail, much shorter than the 20 - 25 residues in a flexible initiation region usually required for efficient insertion and engagement by the ATPase hexamer ^3,29^. This substrate was previously demonstrated to be recruited to the proteasome and slowly deubiquitinated, yet not degraded ^29^. Crosslinking with Rpt1-P1L-Bpa/Rpn11^AXA^ proteasomes and a standard 35 amino acid tail substrate resulted in the expected smeared supershift of the Rpt1 band, while incubation with the single amino acid tail substrate showed only Rpt-Rpt crosslinking (Fig. 2C). Taken together, these results indicate that crosslinking of Bpa-containing 26S proteasomes occurs robustly with substrates that are engaged and translocated by the ATPase motor, while excluding proteins that are degradation-incompetent or bind only transiently to the exterior of the proteasome.

### Rpt1-Bpa expression in yeast cells results in rapid crosslinking to native substrates

After verifying *in vitro* that Bpa incorporation into Rpt1 or Rpt2 results in functional 26S proteasomes and specific crosslinks to engaged substrates in a UV-dependent manner, we aimed to implement this approach in live cells. The 26S proteasome is essential to eukaryotic cells, such that making all proteasome complexes crosslinkable and potentially stall on substrates could lead to considerable inhibition of the UPS and undesired responses that may skew our analyses of trapped substrates. We therefore decided to keep all wild-type proteasome subunits in the cell and express an additional copy of Bpa-containing Rpt1 or Rpt2 with a N-terminal His_6_-HA-tag. Incorporation of these copies results in a proteasome subpopulation that is crosslinkable and tagged for affinity purification and Western blotting, while wild-type proteasomes maintain normal UPS function.

Using galactose-inducible copies of His_6_-HA-tagged Rpt1 or Rpt2 with an amber stop codon in the pore-1 loop Tyr position for Bpa placement, together with the tRNA^Bpa^/tRNA synthetase pair for genetic code expansion, we verified the incorporation of these subunits into proteasome holoenzymes in yeast *S. cerevisiae* (Supp. Fig. 6A). Endogenous proteasomes were affinity purified via a FLAG tag on PRE1 (20S CP subunit β4), and Western blotting against the HA tag revealed that Rpt1-PL1-Bpa efficiently assembled into 26S proteasomes (Supp. Fig 6A). Surprisingly, Rpt2-PL1-Bpa failed to incorporate into proteasomes within yeast cells, despite its successful association into functional base subcomplexes in our recombinant *E. coli* expression system (Supp. Fig 5), which highlights the complexity of endogenous proteasome assembly. We hence decided to proceed with Rpt1-PL1-Bpa for our *in vivo* crosslinking experiments.

Galactose induction leads to strong expression of target proteins in yeast, but the diauxic shift from glucose to galactose metabolism also results in major transcriptional changes and cellular stress that ultimately can lead to cell death ^38^. To avoid this level of perturbation that would also affect the proteasome substrate pool, we orthogonally expressed Rpt1-PL1-Bpa using a system that is controlled by a synthetic hybrid hormone receptor and responds to human progesterone ^39^ (Supp. Fig. 6B).

Yeast cells transformed with the tRNA^Bpa^/tRNA synthetase pair were grown in selection media in the presence of Bpa, progesterone, or both to assess any growth defects caused by the addition of either compound or by the expression of an extra Rpt1 copy. After confirming no growth defects (Supp. Fig 6C), cells were grown in the presence of 2 mM Bpa and 50 nm progesterone for all subsequent experiments. To measure *in vivo* crosslinking, cells were harvested in late log-phase, washed, re-suspended in PBS buffer, and split into two samples, with one kept in the dark and the other placed under a 365 nm UV lamp for 30 minutes at 22 °C (Fig. 3A). After cell harvesting, lysis, and SDS-PAGE, the anti-HA Western blotting revealed a UV-dependent disappearance of the band for HA-His_6_-tagged Rpt1 and the corresponding appearance of a higher molecular weight smear, similar to the results observed after *in vitro* crosslinking with reconstituted 26S proteasomes and a ubiquitinated model substrate (Fig. 3B). For our initial *in vitro* crosslinking experiments, we used an incandescent UV light bulb (365 nm, 100 W) that worked well for small-scale samples but had a narrow illumination area for cross-linking. The incandescent bulb also generated heat and consequently lead to sample evaporation. We therefore switched to an LED lamp of similar wattage that did not heat up samples, yet produced even slightly more efficient crosslinking (Fig. 3B, right lane) and was thus used for all subsequent experiments.

**Figure 3.**
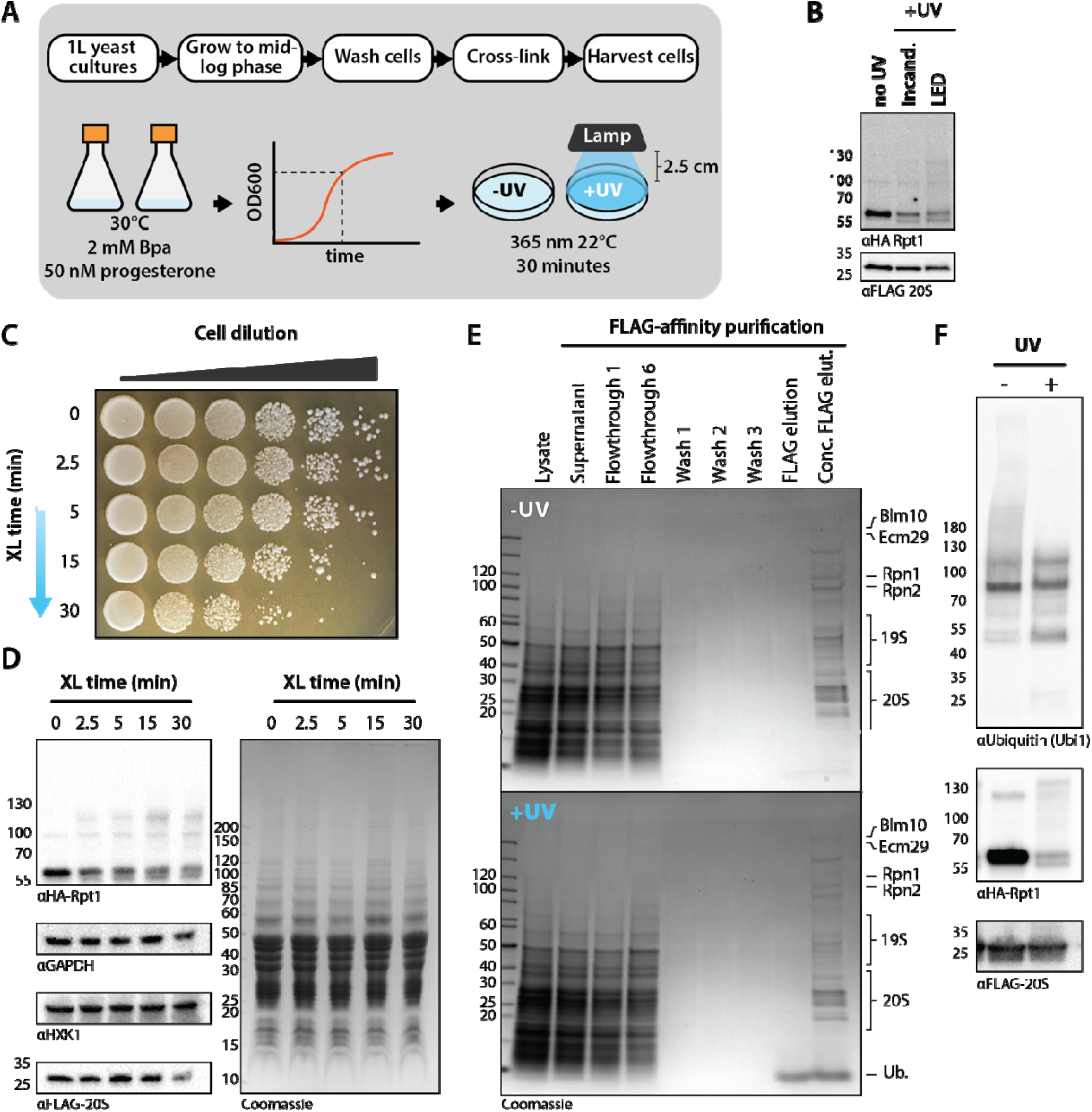
Rpt1-Bpa expresses, incorporates into functional 26S proteasomes, and crosslinks *in vivo*. **A.** Experimental setup for the expression of Bpa-Rpt1 and cross-linking in yeast cells. Crosslinking was carried out with either an incandescent or LED lamp (365 nm) at 22 °C. **B.** Anti-HA Western blot of yeast lysate after crosslinking shows a UV-dependent shift of Rpt1 to a smear as seen in *in vitro* crosslinking with reconstituted proteasomes. **C.** Yeast spot assay confirms that cells remain viable after exposure to 365 nm light. After 30 minutes of exposure, both cell viability and extent of crosslinking decrease. **D.** Western blots for HA-Rpt1, GAPDH, HXK1, and FLAG-20S CP in yeast cell lysates demonstrate that crosslinking is optimal after 15 minutes of UV exposure, with a decrease at 30 minutes. **E.** Anti-FLAG affinity purification of 26S proteasomes following crosslinking demonstrates that the 26S holoenzyme is not disassembled upon crosslinking. UV-exposed cells display an additional band corresponding to free ubiquitin. **F.** Top: Anti-ubiquitin Western blotting of the FLAG elution sample shows that poly-ubiquitinated species co-purify with 26S proteasomes despite extensive wash steps (top, anti-ubiquitin), and this presence of poly-ubiquitin is significantly reduced after crosslinking. Middle: Anti-HA-Rpt1 Western blotting reveals that purified 26S proteasomes elute with cross-linked substrates. Western blotting against the FLAG-tagged 20S CP (bottom) was used as a control for similar proteasome purification and gel-loading.

Concerns about UV-crosslinking include potential light-induced stress and DNA damage, which might affect the proteasome-degraded proteome. Previous mass spectrometry-based studies have shown minimal changes in protein expression following 365 nm light exposure, and this wavelength is extensively used in biological applications ^40^. However, to determine a suitable crosslinking duration that causes minimal cellular interference, we carried out a time course experiment measuring the extent of crosslinking and cell viability after UV exposure. Resuspended cells were exposed to 365 nm light for 2.5, 5, 15, or 30 min, before samples were taken for SDS-PAGE and Western-blot analyses or a spotting assay to test for cell viability. Although cells from all samples were viable, there was a ∼ 5-fold reduction in the number of live cells when increasing exposure from 5 to 15 min and another ∼ 10-fold reduction for extending from 15 to 30 min (Fig. 3C). The Western blotting against HA-Rpt1-BL1-Bpa revealed that crosslinking to substrates occurs already within 2.5 min of UV exposure, further increases up to 15 min, and then slightly declines at 30 min (Fig. 3D), possibly due to cell death or damage-response mechanisms. For our proteasome substrate-profiling experiments, we therefore opted to use a crosslinking time of 15 min.

The timing and mechanisms involved in resolving a proteasome-substrate stall remain largely unknown. It is possible that irreversibly stalled proteasomes, in our case induced through substrate crosslinking, are removed by autophagy or get otherwise disassembled on the timescale of our experiments. To measure the stability of stalled 26S proteasome with crosslinked substrates, we affinity-purified proteasome holoenzymes after UV exposure using the FLAG tag on the 20S CP. Coomassie stained SDS-PAGE gels confirmed that the endogenous proteasomes remained intact, with all intrinsic subunits and even sub-stoichiometric binders like Ecm29 or Blm10 present in the preparation (Fig. 3E, last lane). Importantly, the Western blot against HA-tagged Rpt1-PL1-Bpa revealed a clear high molecular weight smear (Fig. 3F, middle), indicating that the stalled 19S RP with bound substrate remains associated with the 20S CP. When blotting against ubiquitin, we observed a significant amount of higher molecular weight species for the sample that was kept in the dark, even after extensive washing during the proteasome affinity purification (Fig. 3F, top). Surprisingly, the UV-illuminated proteasome sample showed a much lower signal for such large ubiquitin adducts. This difference may indicate potential changes in regulatory ubiquitin modifications of proteasomal subunits themselves after the ATPase motor stalled on a substrate or that ubiquitin chains are more efficiently removed from stalled substrates than from proteins that transiently bind actively degrading proteasomes.

In summary, our proof-of-principle experiments confirmed that Rpt1-PL1-Bpa can be efficiently incorporated into endogenous yeast 26S proteasomes and that UV illumination results in the crosslinking to native substrates, without causing the dissociating or degradation of the stalled proteasome-substrate complexes. However, the presence of co-purifying poly-ubiquitinated species in our uncrosslinked samples (Fig. 3F, left lane) suggests that a simple affinity purification may not be sufficient to remove all non-engaged proteasome binders prior to the mass-spectrometric identification of true substrates.

### Isolating Rpt1-crosslinked substrates under denaturing conditions

Mass spectrometry is an inherently concentration-dependent technique. To increase the signal for Rpt1-crosslinked substrates and eliminate unengaged ubiquitinated proteins or proteasome binders as much as possible, we removed all proteasomal subunits by performing immobilized metal affinity chromatography (IMAC) under denaturing conditions, using the HA-His_6_-tag on Rpt1. SDS-PAGE analyses of test purifications for uncrosslinked proteasome samples in the presence of GdmCl showed a relatively clean band corresponding to Rpt1-PL1-Bpa in the elution from Ni-NTA resin (Supp. Fig. 7). This distinct band disappeared and a higher molecular weight smear could be seen on gels for the denaturing purification of the crosslinked proteasome, consistent with a successful isolation of Rpt1-crosslinked endogenous substrates. Having confirmed this enrichment of Rpt1-substrate adducts, we repeated the UV crosslinking and denaturing purification, followed by analyses with high-resolution mass spectrometry. The yeast proteome contains numerous histidine-rich proteins that appear as contaminants from IMAC purifications along with non-specific interactors. We therefore performed the same mass-spec analysis for the purified uncrosslinked proteasomes to allow a label-free quantification and comparison to background binding of the Ni-NTA resin. Previous work to profile the yeast ubiquitinome by Ni-NTA pull-down of His_6_-tagged ubiquitin followed by mass spectrometry identified 17 histidine-rich contaminant proteins ^27^, and we identified six additional Ni-NTA-binding proteins in our data. Given that these proteins consistently bind as contaminants to Ni-NTA resin and are neither enriched nor depleted in a crosslinking-dependent manner, we used them as internal standards. Indeed, we observed that out of the 17 previously identified histidine-rich proteins only one, the transcription factor Pzf1, showed a significant two-fold enrichment in the crosslinked sample compared to the uncrosslinked control (Fig. 4A, Supp. Fig. 10A). Interestingly, while this his-contaminant appeared as a substrate in this log-phase growth data set, in subsequent experiments for cells experiencing ER-stress this protein dropped out of the hit pool, which further validates it as a *bona fide* log-phase substrate (Supp. Fig. 10B).

**Figure 4.**
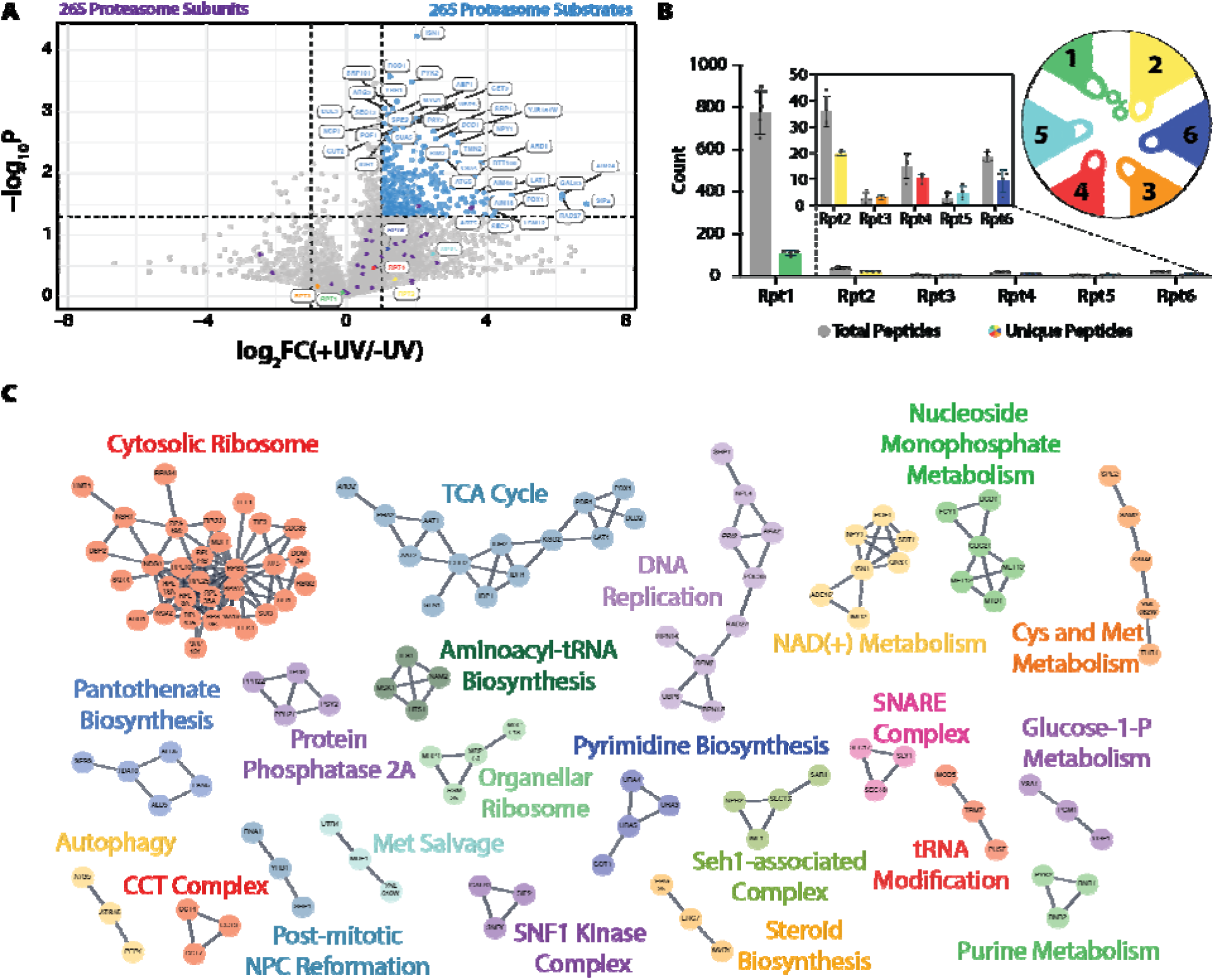
Mass-spectrometry identification of 26S proteasome substrates during log-phase growth. **A.** Volcano plot of proteins identified by mass spectrometry with label-free quantification. Shown in blue are 344 proteins that were at least 2-fold enriched in the crosslinked sample compared to the non-crosslinked control. Most of the 26S proteasome subunits (highlighted in purple) and specifically the Rpt1-Rpt6 motor subunits (shown in various colors) are not significantly enriched. **B.** Mass-spec abundance for Rpt1-Rpt6 motor subunit as measured by counts of peptide-spectral matches. For each Rpt, gray bars show the total number peptide-spectral match counts and colored bars denote the count of unique peptides. The circular diagram depicts the order of Rpt subunits in the ATPase hexamer and the orientation of pore-1 loops within the central channel, pointing toward the clockwise-next Rpt, which explains the higher abundance of Rpt2-derived peptides upon Rpt1-P1L-Bpa crosslinking. **C.** Top clusters of identified proteasome substrates by physical and functional interactions. Colors and labels represent their functional enrichment annotations.

As discussed above, our initial *in vitro* crosslinking experiments with reconstituted proteasomes showed a band on SDS-PAGE gels at approximately the size of an Rpt-Rpt dimer. We therefore decided to analyze our mass-spec data for the *in vivo* crosslinked proteasome regarding Rpt1-bound subunits of the ATPase hexamer. Quantifying the number of total and unique spectra originating from Rpt1-Rpt6 confirmed Rpt1 as by far the most abundant ATPase subunit, with >20 times more peptides than the second most abundant subunit, Rpt2 (Fig. 4B). This result is expected for our denaturing purification of His_6_-tagged Rpt1, which disrupts the subunit interactions of the normally tightly bound ATPase hexamer. Our observations of Rpt2 as the second most abundant subunit and Rpt3 as the least represented Rpt are consistent with their positions in the ATPase hexamer, the orientation of pore-1 loops in the central channel, and the consequent propensities of crosslinking to the Bpa in Rpt1. Rpt1 points with its pore-1 loop toward the clockwise-neighboring Rpt2, leading to potential crosslinking if there is no substrate polypeptide to be targeted (Fig. 4B, inset).

Modifying our database search to consider variable modifications on Tyr residues allowed us to identify spectra corresponding to the uncrosslinked Bpa-containing pore-1 loop peptide (Supp. Fig. 8A). Interestingly, comparing the data for crosslinked and uncrosslinked proteasomes revealed that there was a significant reduction of peptide-spectral matches for this Bpa-containing peptide in the crosslinked samples. In contrast, the pore-2 loop peptide did not exhibit any significant reduction in number of features across the two conditions (Supp. Fig. 8B). To further investigate this, we plotted the fold-enrichment of each peptide identified for both +UV and -UV conditions along the sequence of Rpt1. This analysis indeed showed that peptides corresponding to the Bpa-containing pore-1 loop were depleted in the cross-linked samples (Supp. Fig. 8B). We attributed this decrease to the UV-dependent crosslinking and resulting convolution with spectra of substrate peptides. Attempts to perform a similar analysis for other Rpt subunits were unsuccessful due to insufficient peptide coverage for those non-enriched subunits.

Non-proteasomal proteins identified in the mass-spec data for the crosslinked proteasome sample were considered as putative substrates if they were enriched at least two-fold compared to the uncrosslinked control across four biological replicates. By this criterion, our analyses identified 343 potential proteasome substrates (Fig. 4A). Proteins were clustered by their high-confidence functional and physical interactions, and these clusters were analyzed for enrichment in functions (Figure 4C) ^41^. This clustering analysis demonstrated that the pool of proteasome substrates in actively dividing cells is indeed highly diverse. Our data showed 47 clusters of putative proteasome substrates with diverse cellular functions that correlated well with processes expected for log-phase growth. By far the largest pool of substrates (35 proteins) represents proteins involved in cytoplasmic translation, including 10 ribosomal proteins. Ribosomal proteins are synthesized in excess and their subsequent reduction in copy number is well-documented as a proteasome-dependent process ^42^. In addition to translation machinery, we also saw distinct functional clusters for DNA replication, central carbon metabolism, and the biosynthesis of nucleotides and essential cofactors, such as NAD and pantothenate. These clusters are in good agreement with processes that support log-phase cell growth and hence also depend on the regulation through proteasomal degradation.

### Proteasome crosslinking can profile changes in the substrate pool

To verify the ability of our *in situ* substrate crosslinking approach to profile changes in proteasome substrate pools, we induced protein misfolding stress at the endoplasmic reticulum (ER) via treatment with the drug tunicamycin. Tunicamycin is a nucleoside analog that competitively inhibits the UDP-N-acetyl-glucosamine-1-P transferase (GPT) enzyme Alg7 (Fig. 5A). On the cytosolic face of the ER, Alg7 carries out the transfer of GlcNAc-1-P from UDP-GlcNAc to a dolichyl-phosphate lipid (Dol-P), which represents the first step in protein glycosylation at the ER. Other sugars are then added to the Dol-P-bound glycan before its ‘en bloc’ transfer to proteins during their translation into the ER lumen or membrane ^43^. The addition of tunicamycin hence results in defective glycosylation of ER proteins, activation of the unfolded protein response (UPR), and the retrotranslocation of proteins from the ER to the cytosol for degradation by the 26S proteasome ^44,45^.

**Figure 5.**
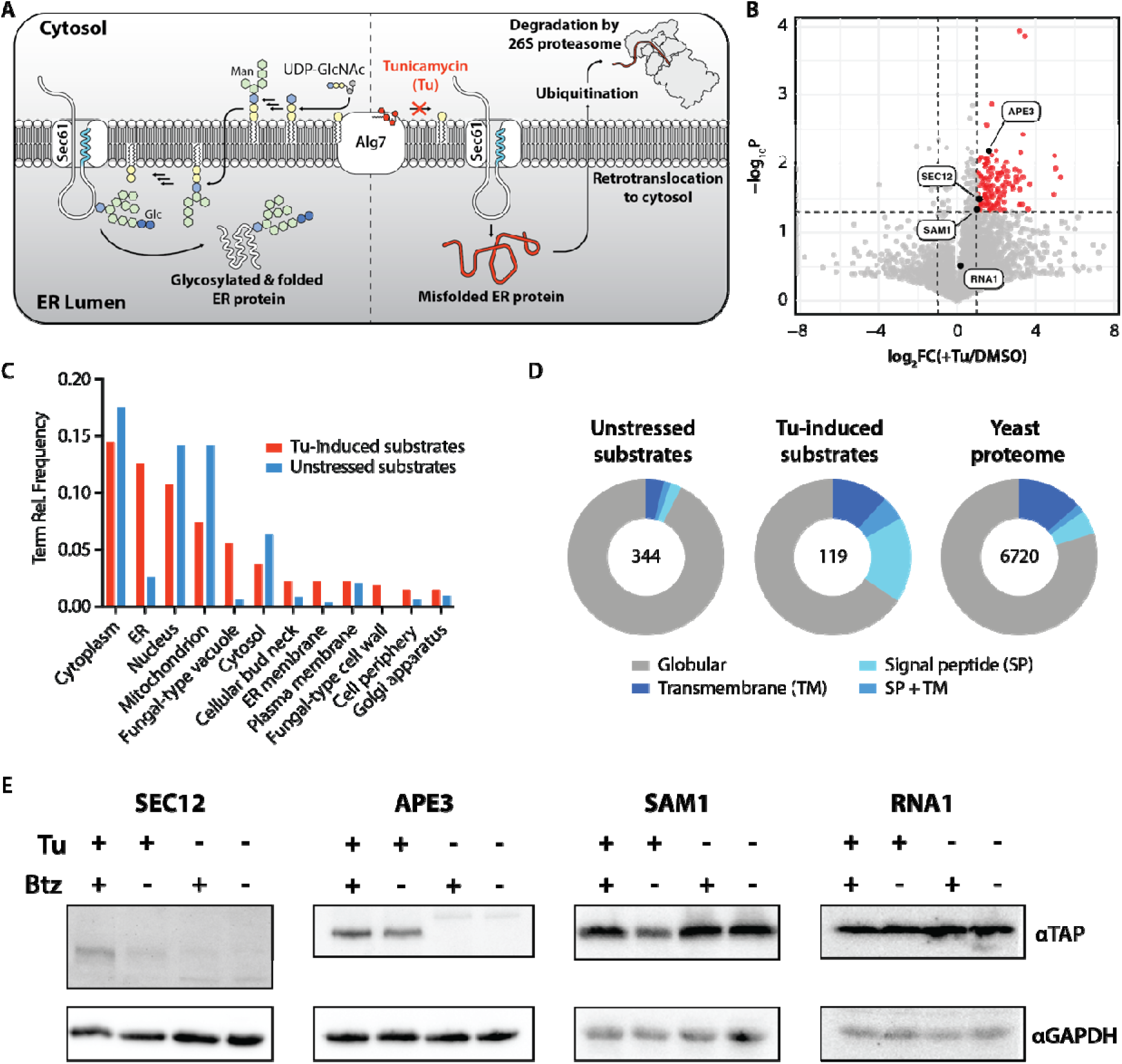
Live-cell crosslinking reveals changes in 26S proteasome substrates induced by endoplasmic reticulum stress. **A.** Schematic for the effect of tunicamycin (Tu) on protein glycosylation and folding in the endoplasmic reticulum (ER). **B.** Volcano-plot comparison of proteins identified following denaturing Ni-NTA enrichment from crosslinked cells treated with Tu or DMSO. Shown in orange are 119 proteasome substrates that are at least 2-fold enriched upon cell treatment with Tu. The four proteins used for further validation in panel E are labelled. **C.** Relative abundances of cellular localization terms for all 26S proteasome substrates identified during log-phase growth and after Tu-induced ER stress. **D.** DeepTMHMM prediction of globular, transmembrane, and signal-peptide-containing proteins among the proteasome substrates identified during log-phase growth and after TU-induced stress, in comparison to the total yeast proteome. **E.** Representative Western blots for four different proteins that represent distinct modes of Tu-induced degradation. Sec12 and Ape3 show increased degradation in response to increased expression, with different magnitudes of degradation. Sam1 shows Tu-induced degradation without changes in expression. By contrast, Rna1 was not identified as a Tu-induced substrate and correspondingly shows no increased degradation upon Tu addition.

For inducing ER stress, we added tunicamycin (in DMSO) to the growth medium 2 hours prior to performing the proteasome crosslinking (Supp. Fig. 9A). As a control for a label-free comparison, we performed the same experiments with cells in the presence of just DMSO. The desired effect of tunicamycin on ER-protein maturation and folding was confirmed by testing for UPR activation, for which we extracted RNA from treated cells and verified the splicing of the mRNA for Hac1 by RT-PCR (Supp. Fig. 9B).

From the crosslinking experiments in the presence of tunicamycin we identified 119 proteins that were significantly enriched or only observed upon tunicamycin treatment compared to untreated cells for all biological replicates (Fig. 5B). To assess whether the substrates from tunicamycin-stressed cells show an enrichment of ER-resident proteins and proteins of the secretory pathway, we used cellular localization annotations from the Saccharomyces Genome Database (yeastgenome.org), mapped the proteins from both the stressed and unstressed cell datasets to their annotated localizations, and plotted the relative frequencies of localization terms shared across the datasets. As expected, terms corresponding to the ER, ER membrane, vacuole, and cell wall were highly represented in our tunicamycin-induced substrates as compared to substrates trapped in unstressed cells (Fig. 5C). We noticed that substrates with annotations for cytosolic or nuclear localization were not reciprocally reduced to a similar extent as membrane-associated substrates increased in the tunicamycin-treated sample. This suggests that the cellular proteasome pool has sufficient capacity to maintain a high degradation level for normal, non-tunicamycin-induced substrates while handling the additional load of misfolded or mistargeted proteins originating from the ER stress.

Given that the glycosylation inhibition by tunicamycin results in ER stress due to faulty protein processing, we expected that substrates containing transmembrane domains and signal peptides for transport into the ER would be highly represented in our tunicamycin dataset. We therefore employed the Deep Learning model DeepTMHMM to computationally predict the membrane topology of the protein substrates identified from unstressed and tunicamycin-stressed cells ^46^. As a comparison, we also ran this analysis on the entire yeast proteome. DeepTMHMM predicted that 80% of the 6720 submitted yeast proteome sequences are globular and soluble, 15% contain transmembrane domains with or without signal peptides, and ∼ 5% were detected to contain signal peptides for import into the ER (Fig. 5D). 92% of the substrates identified from unstressed cells during log-phase growth were globular proteins and depleted of transmembrane domains or signal peptides when compared to the overall yeast proteome. Only ∼ 5% of these substrates contained transmembrane domains and 2% had a signal peptide. Of the trapped substrates enriched upon tunicamycin treatment, 17% were transmembrane domain-containing and nearly 18% included a signal peptide, with the fraction of predicted globular proteins dropping to 66%. These results confirmed that the most prominent actively degraded substrates were distinct for the two experimental conditions. Exposure to ER stress led to the strongly increased degradation of substrates with typical characteristics for ER-resident or secretory proteins.

To further confirm that proteins crosslinked to the proteasome upon tunicamycin stress indeed represented substrates being degraded in a proteasome-dependent manner, we employed a yeast library of TAP-tagged ORFs and measured the tunicamycin-induced degradation for a subset of protein hits. By Western blotting against the TAP tag, we monitored differences in the abundance of these proteins for cells grown in the absence or presence of tunicamycin (Tu) and the proteasome inhibitor bortezomib (Btz). We measured the proteasome-dependent degradation of SEC12, APE3, and SAM1 upon addition of tunicamycin, which showed various levels of turnover (Fig. 5E). SEC12 is a membrane-localized ER guanine exchange factor (GEF) protein, APE 3 is a vacuolar aminopeptidase, and SAM1 is a metabolic enzyme that catalyzes the biosynthesis of S-adenosylmethionine. Curiously, in the presence of tunicamycin and bortezomib, lower molecular weight species of both SEC12 and APE3 appear to increase in abundance. This shift to a lower apparent molecular weight is likely due to defective glycosylation, which is resolved via degradation by the 26S proteasome. As a control, we performed the same experiment on RNA1, which was identified as a log-phase substrate but not during tunicamycin-induced stress, and it indeed did not show changes in degradation upon tunicamycin addition (Fig. 5E). Taken together, these results further demonstrate that the putative substrates identified with our approach represent a snapshot of the proteasome-engaged substrate pool at the time of crosslinking and not just changes in protein expression upon ER stress.

## Discussion

The 26S proteasome functions as an essential hub for regulated protein degradation and is therefore an important drug target for the treatment of various human diseases, ranging from cancer to neurodegeneration. The advent of induced protein degradation through proteolysis targeting chimeras (PROTACs) or molecular glues further emphasizes the proteasome’s role as a viable therapeutic modality and highlights the importance of understanding its cellular substrate pool. Almost every process in the eukaryotic cell is at some point regulated by the ubiquitin-proteasome system, yet the identity of its substrates and especially how the primary targets for proteasomal degradation change in response to altered cellular needs or stresses remains largely elusive. So far there has been no robust strategy for elucidating the pool of proteins that get degraded by the 26S proteasome in an ATP-dependent and ubiquitin-mediated or ubiquitin-independent manner. Proteomic studies of changes in protein abundances after the inhibition of certain processes or pathways are generally limited by various feedback mechanisms and responses in protein expression, trafficking, or turnover. Similarly, analyses of the ubiquitinated proteome cannot provide a direct picture of proteasomal substrates, as not all proteins modified with ubiquitin chains of proteasome-targeting linkage types indeed end up getting degraded. On the other hand, mass-spec analyses of potential proteasomal cleavage products found in proximity to the 20S CP in lysates have the disadvantage of limited specificity, because they may report on hydrolysis by various capped proteasome complexes and secondary cleavage events of the uncapped 20S CP.

In this work, we demonstrate that capturing proteins within the central channel of the 19S regulatory particle is a powerful approach for profiling committed proteasomal substrates on their way to degradation. Structure-based mutagenesis and genetic code expansion allowed us to introduce Bpa into the pore-1 loop of Rpt1 for specific photo-induced crosslinking of substrates, despite the high architectural complexity of the 26S proteasome. This strategy is viable for substrate trapping in live yeast cells by using the orthogonal expression of an Bpa-containing additional copy of Rpt1, which efficiently incorporates into the 26S holoenzyme and generates a subpopulation of crosslinkable proteasomes that does not interfere with normal UPS function. Purification of Rpt1-substrate adducts under denaturing conditions enabled us to eliminate other proteasome interactors or transiently bound proteins and provided a snapshot of the very broad substrate pool in unstressed cells during log-phase growth. As a proof-of-concept we utilized this crosslinking mass-spec approach to analyze the shift in the proteasomal substrate pool after exposure to tunicamycin-induced ER stress. The resulting change in trapped substrates, most notably the enrichment of ER-resident or secretory proteins and proteins with signal peptides or transmembrane domains, validates our strategy as a powerful tool for interrogating differences in proteasome substrate preferences depending on growth conditions or stress responses. We envision this specific crosslinking approach as a groundbreaking method for elucidating how the ubiquitin-proteasome system regulates numerous vital processes and shapes the eukaryotic proteome based on cellular needs.

## Resource availability

### Lead contact

Further information and requests for resources and reagents should be directed to and will be fulfilled by the lead contact, Andreas Martin.

### Materials and data availability

Data generated or analyzed during this study are included in the manuscript and the Supporting Information. The raw mass spectrometry data were deposited into the ProteomeXchange Consortium via the PRIDE partner repository under the dataset identifier PXD080919 for proteasome crosslinking during log-phase growth and PXD080917 for proteasome crosslinking under ER stress. This paper does not report original code. All constructs are available from the lead contact upon request and completion of a Material Transfer Agreement.

## Acknowledgments

The authors thank all members of the Martin Lab for valuable discussions. We also thank Lei Wang (UCSF) for kindly providing the pSNR-TyrRS plasmid, Joseph Loebbel and Nick Ingolia (UC Berkeley) for reagents and guidance in orthogonal expression in yeast, and Robert Maxwell and the Vincent J. Coates Proteomics/Mass Spectrometry Laboratory Core Facility in the QB3 Institute for the development of methods necessary for the success of this work.

## Funding

This research was funded by a Gilliam Fellowship from the Howard Hughes Medical Institute (S.Y.R.), the Howard Hughes Medical Institute (S.Y.R. and A.M.), and by the US National Institutes of Health (R01-GM094497 to A.M.).

## Author Contributions

Conceptualization, S.Y.R. and A.M.; Methodology, S.Y.R. and A.M.; Investigation, S.Y.R.; Writing – Original Draft, S.Y.R. and A.M.; Writing – Review & Editing, S.Y.R. and A.M.; Funding Acquisition, A.M.; Supervision, A.M.

## Declaration of Interests

The authors declare no competing interests.

## Methods

### Construction of plasmids for Bpa incorporation in *E. coli* and yeast

For Bpa incorporation in our recombinant proteasome expression system it was necessary to clone the tRNA and tRNA synthetase pair from the pEVOL-pBpF plasmid originally produced by the Schultz lab ^47^ into a pULTRA plasmid backbone for IPTG-inducible expression of the Bpa-incorporating tRNA/synthetase pair. To do this, the pULTRA backbone of the pAzFRS.2.t1 plasmid, the tRNA, and the *M. jannaschii* tRNA synthetase from the pEVOL plasmid were PCR-amplified to add homologous overhangs for subsequent Gibson assembly in the pULTRA backbone downstream of a tac promoter.

For Bpa incorporation in live cells, the pSNR-EcTyrRS-tRNA plasmid that contains the amber-decoding tRNA_CUA_ and a wild-type *E. coli* tyrosine tRNA synthetase was mutated such that the synthetase accommodates Bpa. The plasmid was cut with the restriction enzymes BamHI and SpeI to drop out a segment of the Ec-TyrRS gene, and a gBlock (IDT) of the same fragment bearing mutations Y37G, D182G, and L186A was Gibson assembled into the cut backbone to produce the pSNR-BPA-RS plasmid.

### Protein expression and purification

Purification of Bpa-containing base subcomplexes was carried out as previously described ^30,48^. In short, stocks of *E. coli* BL21(DE3) cells containing the pET plasmid (coding for base subunits Rpn1, Rpn2, and Rpn13) and pACYC plasmid (coding for assembly chaperones) were sequentially transformed by electroporation, first with the pULTRA-Bpa2 plasmid for UAA incorporation and then the pCOLA plasmid (coding for Rpt1-Rpt6) for the expression of amber codon-containing motor subunits. Quadruple transformants were plated on LB plates supplemented with ampicillin, kanamycin, chloramphenicol, and spectinomycin. For each variant, a single colony was picked and an overnight culture was grown in dYT media under quadruple selection. With the overnight culture, 6 liters of dYT media were started and grown at 37 °C to an OD_600_ of 0.6. Cells were harvested by centrifugation and resuspended in 1 L unbuffered Terrific Broth (TB) media for six-fold cell concentration. Bpa was dissolved in NaOH and added to the TB to a final concentration of 2 mM. Protein expression was induced by the addition of 1 mM IPTG and allowed to progress for 5 hours at 30 °C and then 16 °C overnight. Cells were harvested by centrifugation, and pellets were either frozen or carried forward to purification. Cell pellets were resuspended in NiA buffer (60 mM HEPES, 100 mM NaCl, 100 mM KCl, 10 mM MgCl_2_, 20 mM imidazole, 5% glycerol, pH 7.6) supplemented with protease inhibitors and lysed by sonication. The cleared lysate was batch-bound for 1 hour to Ni-NTA agarose to immobilize the 6xHis tag on Rpt1, beads were washed, and proteins were eluted with NiB buffer (60 mM HEPES, 100 mM NaCl, 100 mM KCl, 10 mM MgCl_2_, 250 mM imidazole, 5% glycerol, pH 7.6). The Ni-NTA elution sample was then flowed several times over a column containing anti-FLAG IgG resin to bind the FLAG-tag on Rpt3, washed, and eluted using excess FLAG peptide. FLAG elutions were concentrated to 500 μL and further cleaned up by size-exclusion chromatography in GF buffer (30 mM HEPES, 50 mM NaCl, 50 mM KCl, 10 mM MgCl_2_, 5% glycerol, pH 7.6) using a pre-equilibrated Superose 6 Increase column. Fractions containing the proteasome base subcomplex were concentrated using a 30K MWCO spin concentrator, the concentration was determined by Bradford method, and aliquots were flash frozen.

Purification of additional proteasome components have been previously described ^30^. In short, the lid subcomplex was expressed in *E. coli* BL21(DE3) that contained the three plasmids coding for lid subunits and the Hsp70 chaperone. The two-step affinity purification first employed a Ni-NTA resin to bind the 6xHis tag on Rpn12, followed by a maltose resin to immobilize a MBP-Rpn6 fusion. The lid subcomplex was further purified by size-exclusion chromatography on a pre-equilibrated Superose 6 Increase column. Purification of the *S.c.* Rpn10 was conducted as previously described ^16^. Briefly, *E. coli* BL21(DE3) cells were transformed with pACYCDuet-1 containing 6xHis-scRpn10 and grown in DYT media at 37 °C until OD_600_ of 0.6-0.8. Protein expression was induced with 0.5 mM IPTG at 37 °C for 4 hours. Cells were harvested and resuspended in lysis buffer (60 mM HEPES, pH 7.6, 100 mM NaCl, 100 mM KCl, 20 mM imidazole, 10% glycerol) containing 2 mg/mL lysozyme, benzonase, and protease inhibitors (aprotinin, pepstatin, leupeptin and PMSF). The resuspended pellet was stored at -80°C until purification. Cells were lysed by sonication, and the clarified lysate was bound in batch to 5 ml of Ni-NTA affinity resin (Thermo Fisher). The resin was washed with lysis buffer and eluted with lysis buffer + 250 mM imidazole. 2 mM DTT was added directly to the elution and the protein was stored overnight on ice. The next day the protein was purified using a Superdex 75 16/60 column (Cytiva) pre-equilibrated with lysis buffer. Peak fractions were concentrated, flash frozen and stored at -80°C.

Endogenous 20S core peptidase was purified from a yeast strain in which a FLAG tag was genomically added to the PRE1 gene, which codes the □4 subunit. Yeast cells were grown to saturation in YPD media, pelleted, resuspended in GF buffer, flash frozen (popcorned into liquid nitrogen), and then lysed using a cryogenic mill. The yeast lysate powder was resuspended in GF buffer and run over an anti-FLAG IgG resin, washed with a high salt buffer to separate endogenous 19S RP, and 20S CP was eluted with excess FLAG peptide. The 20S CP was then further cleaned up and buffer exchanged by size-exclusion chromatography on a pre-equilibrated Superose 6 Increase column. Concentrations were determined by absorbance at 280 nm and aliquots were flash frozen.

To purify the titin-I27(V13P/V15P)-cyclinB model substrates, a starter culture of *E. coli* BL21(DE3) cells carrying the expression construct for the substrates was made and allowed to grow overnight. The following day, starter cultures were centrifuged, cells were washed with fresh media, and diluted into M9 Minimal media with added M9 salts and kanamycin to a starting OD_600_ of 0.05. Cells were grown at 30 °C to an OD_600_ of 0.7, and protein expression was induced by the addition of IPTG to a final concentration of 0.5 mM. Expression was carried out for 3 hours at 30 °C. Cells were pelleted, resuspended in Chitin Buffer (60 mM HEPES, 150 mM NaCl, 2 mM EDTA, 5% glycerol, pH 7.6, supplemented with proteasome inhibitors: 1 mM AEBSF and 1 μg/mL each of aprotinin, pepstatin, and leupeptin), and flash frozen. For purification, the pellet was thawed and lysed by sonication. The lysate was clarified by centrifugation, and the supernatant was batch-bound for 1 hour to chitin resin pre-equilibrated with Chitin Buffer. The resin-lysate mixture was transferred to a gravity column, washed, and incubated with elution buffer (30 mM HEPES, 100 mM NaCl, 2 mM EDTA, 5% glycerol, pH 8.5) overnight to cleave the chitin-binding domain. Concentrations of the eluted titin substrates were determined by absorbance at 280 nm, and aliquots were flash frozen.

### Substrate ubiquitination and labelling

Purified titin substrates were labelled with a fluorescein peptide using sortase in a 0.5 mL reaction, containing 20 μM SrtA, 10 mM CaCl_2_, 0.5 mM fluorescein-amidite (FAM)-labeled peptide, 100 μM substrate, and 1 mM DTT in GF buffer. The reaction was incubated in the dark for 1 hour at room temperature, then diluted with two volumes of labelling buffer (30 mM HEPES, 300 mM NaCl, 20 mM imidazole, 5% glycerol, pH 7.6), loaded onto a 1 mL Ni-NTA cartridge to remove the His-tagged SrtA enzyme, washed with binding buffer, and eluted with elution buffer (30 mM HEPES, 300 mM NaCl, 500 mM imidazole, 5% glycerol, pH 7.6). The elution was concentrated to 500 μL and run on a pre-equilibrated Superdex 75 Increase 10/300 size-exclusion column for further cleanup. Fractions corresponding to labelled substrate were pooled, the concentration was determined by absorbance, and aliquots were flash frozen.

Prior to degradation experiments, labelled substrates were ubiquitinated in a 200 μL reconstituted ubiquitination mixture. The mixture was composed of 2.5 μM mouse E1 ubiquitin-activating enzyme, 2.5 μM Ubch7 E2 ubiquitin conjugating enzyme, 5 μM Rsp5 E3 ubiquitin ligase, 50 μM ubiquitin, and 10 μM substrate. The reaction was started by the addition of ATP to 10 mM final concentration and allowed to run for 45 minutes. Ubiquitinated substrates were kept on ice and used in assays on the same day.

### 26S proteasome reconstitution and degradation assay

Multiple-turnover degradation was monitored by decrease in fluorescence anisotropy of FAM-labelled substrate in a Synergy Neo2 plate reader (Biotek) equipped with a filter cube for excitation at 485 nm (20 nm bandwidth) and emission at 528 nm (20 nm bandwidth). Assays were carried out with final concentrations of 50 nM 20S CP, 400 nM wild-type or Bpa-containing base, 600 nM lid, 750 nM Rpn10, and 2 μM ubiquitinated FAM-substrate with 1X ATP-regenerating mixture (0.03 mg/mL creatine kinase and 16 mM creatine phosphate). To prepare the reactions, separate 2X enzyme and 2X substrate mixtures were set up in PCR strips and then mixed to initiate degradation. The 2X enzyme contained all proteasome components in GF buffer, supplemented with 0.5 mM ATP, 0.5 mg/mL BSA, and 0.5 mM TCEP. The 2X substrate mix contained the ubiquitinated FAM-labelled substrate in GF buffer with ATP, BSA, and TCEP. Both reaction mixes were incubated on ice for 15 minutes to allow the 26S proteasome to fully reconstitute. Prior to starting the degradation reactions, the 2X enzyme, 2X substrate, and 384-well flat bottom microplate were incubated at 30 °C for 5 minutes. A multichannel pipette was used to mix the substrate and enzyme mixes to 1X, 10 μL aliquots were quickly loaded onto the plate, and anisotropy measurements were initiated. Anisotropy traces were plotted and analyzed using Prism 10 (GraphPad Software).

### *In vitro* crosslinking

For substrate crosslinking to reconstituted proteasomes *in vitro*, the final concentrations were 600 nM 20S CP, 400 nM Bpa-containing base variant, 600 nM lid, 750 nM Rpn10, and 2 μM substrate. For initial crosslinking experiments the concentrations were adjusted to avoid having unincorporated crosslinkable base subcomplex. 2X concentrated proteasome samples were reconstituted with each tested base variant (Rpt1-P1L or Rpt2-P1L) and lid containing wild-type Rpn11 or Rpn11_AXA_. These proteasome samples were incubated on ice for 15 minutes, and crosslinking reactions were started by mixing with a 2X substrate sample. The uncapped reaction was left on ice and placed under a 365 nm lamp for 15 minutes. Reactions were then quenched by the addition of 4X SDS-PAGE sample buffer and heating to 95 °C. Samples were run on a 5-10% gradient Tris-Glycine PAGE gel and imaged for in-gel FAM fluorescence or transferred onto a PVDF membrane that was subsequently blocked, blotted with an anti-Rpt1 or anti-Rpt2 antibody, incubated with a HRP secondary antibody, and imaged using chemiluminescence on a Bio-Rad ChemiDoc MP. For crosslinking experiments with different substrate variants (with or without an unstructured tail for initiation), the experimental setup was identical except for concentrations, which were the same as in the degradation assays: 50 nM 20S CP, 400 nM wild-type or Bpa-containing base, 600 nM lid, 750 nM Rpn10, and 2 μM ubiquitinated FAM-substrate.

### Western Blotting

For SDS-PAGE, all protein samples were diluted with 4X Laemmli Sample Buffer (Bio-Rad) containing fresh β-mercaptoethanol and boiled at 95°C for 5 minutes prior to loading onto a 10% Tris-Glycine gel and running at 180 V. Gel bands were visualized with Coomassie stain. For Western blotting, samples were transferred onto a PVDF membrane using an iBlot2.0 (Thermo Fisher) using the standard transfer protocol, membranes were blocked with 5% milk in TBST and blotted with primary antibody. Blots were developed using an HRP-conjugated secondary antibody, chemiluminescent substrate solution (Perkin Elmer), and visualization with a Bio-Rad ChemiDoc MP.

### Strain construction Yeast transformation

All yeast-strain construction was carried out via the lithium acetate transformation protocol. For the purification of native holoenzyme from yeast following crosslinking, a FLAG tag was added to the C-terminus of PRE1 (□4 subunit of 20S CP) by transformation of wild-type W303 (MATα, can1-100, his3-11, leu2-3,112, lys2, trp1-1, ura3-1) cells with a PCR amplicon containing the PRE1-FLAG fusion sequence with a -His cassette for selection. Subsequent integrations were made into this strain for additional expression of Rpt1 or Rpt2 variants.

### Expression of Rpt1 and Rpt2

#### GAL expression

For gal-inducible expression of Rpt1 or Rpt2, the sequences of Rpt1-Y283TAG and Rpt2-Y256TAG were PCR amplified to add homology to a pYES-HTH plasmid. The linearized pYES backbone and Rpt-containing amplicons were then assembled using a NEB HiFi DNA Assembly Kit (New England Biolabs). Cells containing the pYES plasmids with Rpt1Y283TAG or Rpt2Y253TAG were then transformed with the pSNR-BPA-RS plasmid. To express the Bpa-containing subunits in yeast, strains were grown in complete synthetic media (CSM) -Ura -Trp supplemented with 0.5% glucose in 50 mL cultures overnight, spun down, washed in media containing 0.5% glucose, and then diluted into large cultures with a starting OD of 0.2. Cells were grown until an OD of 1, and 20% galactose was added to a final concentration of 2%. Cells were grown for 6 hours, harvested by centrifugation, and flash frozen with liquid nitrogen.

#### Progesterone-induced expression

For the orthogonal expression of Rpt1Y283TAG in live yeast, a two-plasmid progesterone expression system was inserted genomically using EasyClone2.0 plasmid backbones. Yeast strains were first transformed to integrate a constitutively expressed Z4PM synthetic receptor and then plated and selected for the hygromycin resistance gene present on the EasyClone backbone. The sequence of HA-6xHis-Rpt1-Y283TAG was cloned downstream of the P(Z4) promoter site and integrated into the yeast cell, which were then selected for resistance by plating on nourseothricin plates. Strains containing genomically-integrated FLAG-tag, the Z4PM synthetic progesterone receptor, and the P(Z4)-HA-6xHis-Rpt1-Y283TAG were sequentially verified by colony PCR, before they were finally transformed with a plasmid containing the tRNA/tRNA synthetase pair for Bpa incorporation and plating on tryptophane drop-out plates.

### Growth curves

A single colony of the yeast strain containing the integrated Rpt1 expression construct and transformed with the pSNR-BPA_RS plasmid was picked and grown in CSM -Trp media overnight in a 10 mL starter culture at 30°C with rotation. After reaching saturation, the OD_600_ of the starter culture was measured and diluted to 0.1 in a new culture, grown for several divisions (∼3 hours) and then diluted to an OD_600_ of 0.05 in media containing 2 mM Bpa, 50 nM progesterone or both. In a 96-well flat bottom plate (Corning Inc.), 100 μL for each condition were added in triplicate with media-only controls for background subtraction. The plate was covered with a Breathe-Easy sealing membrane (Sigma) and read in a Synergy Neo2 plate reader (Biotek) set to 30°C, measuring at 600 nm every 10 minutes for 24 hours. OD_600_ measurements from the plate reader were formatted, exported as a CSV file and analyzed using the Growthcurver R package ^49^, with the media-only samples for background subtraction.

### Yeast crosslinking

The crosslinking-competent yeast cells were streaked on a CSM-Trp plate and grown for 2-3 days. A single colony was picked and grown overnight in 5 mL CSM-Trp media. The 5 mL culture was then used to start several 50 mL overnight cultures. The following day 8 L of CSM-Trp media was made and supplemented with Bpa dissolved in NaOH to a final concentration of 2 mM and 50 nM progesterone from a 1000X stock. The media was inoculated to a starting OD_600_ of 0.1, and cells were grown until OD_600_ 0.7-0.8, which took approximately 13-15 hours. Cells were pelleted by centrifugation, washed once with PBS, resuspended in 400 mL PBS, and divided into 8 sterile 150 x 25 mm tissue culture plates (50 mL of cell suspension per plate). Four plates were left covered at room temperature and four plates each had a 365 nm LED lamp placed on top, which positioned the light source ∼2 cm from the cell suspension. Cells were crosslinked for the desired amount of time and then harvested from the plates, centrifuged, and washed with PBS.

For experiments to determine the optimal crosslinking time, five plates with cell suspensions were left in the dark or exposed to UV light for 2.5, 5, 15, 30 minutes. The cell suspensions were harvested, washed, and 1 mL of suspension for each condition was then ten-fold serially diluted. A spot plate on YPD media was made by pipetting 10 μL of each dilution and incubated at 30°C. For crosslinking experiments to be analyzed by mass spectroscopy, yeast cells were grown as described above. Following the *in vivo* crosslinking, cells were washed once with PBS, resuspended with ice-cold PBS containing protease inhibitors (1 mM AEBSF, 1 ug/mL each of aprotinin, pepstatin, and leupeptin) to a volume of 20 mL, drip frozen in liquid nitrogen, and stored at -80°C until purification.

### Denaturing purification of Rpt1-substrate crosslinks

The yeast pellets were lysed in a SPEX Sample Prep cryogenic mill using a total of 15 3-minute cycle, with 2 minute grinding (15 impact cycles per second) followed by 1 minute of cooling. The yeast lysate powder was resuspended in 60 mL denaturing binding buffer (20 mM Na_2_HPO_4_, 150 mM NaCl, 10 mM imidazole, 8 M GdmCl, pH 7.4) to a final concentration of 6 M GdmCl. Fresh Ni-NTA agarose beads were washed, pre-equilibrated with the binding buffer, and added to the lysate for 1 hour of batch binding. The lysate mixture was added to a fresh disposable chromatography column, and the resin was allowed to settle before snapping the bottom of the column open to allow flow. The resin was washed with denaturing wash buffer 1 (20 mM Na_2_HPO_4_, 300 mM NaCl, 10 mM imidazole, 6 M GdmCl, pH 7.4) followed by a denaturing wash buffer 2 (20 mM Na_2_HPO_4_, 300 mM NaCl, 25 mM imidazole, 6 M GdmCl, pH 7.4). The 6xHis-Rpt1-substrate crosslinks were eluted from the column with denaturing elution buffer (20 mM Na_2_HPO_4_, 300 mM NaCl, 250 mM imidazole, 6 M GdmCl, pH 7.4) and concentrated to ∼1 mL using a 10K MWCO concentrator.

### Mass spectrometry

Purified proteins were methanol precipitated, washed, and resuspended in buffer (50 mM ammonium bicarbonate, 25% acetonitrile, pH 8). Disulfides were reduced with TCEP at a final concentration of 5 mM and then alkylated in 10 mM iodo-acetic acid. Samples were digested with trypsin in 1 mM CaCl_2_. Trypsin-digested peptides were filtered with a 0.22 μm centrifugal filter (Millipore UFC30GVNB) and manually loaded on an in-house prepared fused silica capillary column 250 μm X 35 cm with Kasil outlet end. The column was packed with ReproSil pHoenix C18-3.0 μm high pH resin (Dr. Maisch GmbH). Peptides were eluted at a flow rate of 2 μL/min using a linear gradient of 2–40% buffer B in 140 min (buffer A: 0.1% Triethylamine (TEA) pH 10.0 and 5% acetonitrile in water; buffer B: 0.1% TEA pH 10.0 and 95% acetonitrile in water) with an Agilent 1200 nanoflow LC. Samples were collected every 3 minutes during the gradient using a fraction collector. The 60 fractions were concatenated to 8 fractions, acidified with formic acid to pH 2.0, and reduced in volume to 10 μL.

Digested peptides were analyzed by online capillary nanoLC-MS/MS using a 25 cm reversed phase column and a 10 cm pre-column fabricated in-house (75 µm inner diameter, packed with ReproSil-Gold C18-1.9 μm resin (Dr. Maisch GmbH)) that was equipped with a laser-pulled nanoelectrospray emitter tip. The precolumn used 3.0 μm packing (Dr. Maisch GmbH). Peptides were eluted at a flow rate of 300 nL/min using a linear gradient of 2–40% buffer B in 140 min (buffer A: 0.05% formic acid and 5% acetonitrile in water; buffer B: 0.05% formic acid and 95% acetonitrile in water) in an Thermo Fisher Easy-nLC1200 nanoLC system. Peptides were ionized using a FLEX ion source (Thermo Fisher) using electrospray ionization into a Fusion Lumos Tribrid Orbitrap Mass Spectrometer (Thermo Fisher Scientific). Data were acquired in orbi-trap mode. Instrument method parameters were as follows: MS1 resolution, 120,000 at 200 m/z; scan range, 350−1600 m/z. The top 20 most-abundant ions were subjected to collision-induced dissociation with a normalized collision energy of 35%, activation q 0.25, and precursor isolation width 2 m/z. Dynamic exclusion was enabled with a repeat count of 1, a repeat duration of 30 seconds, and an exclusion duration of 20 seconds.

Datasets in RAW file format were analyzed using FragPipe with a label-free quantification (LFQ) workflow comparing four replicates of +UV to -UV samples. Spectra were matched against a database of the yeast proteome with decoys. Peptide-spectral matching was carried out with MSFragger with a precursor mass tolerance range of -/+20 ppm and a fragment mass tolerance of 20 ppm. Enzymatic cleavage was set to strict tryptic digest at the C-terminal site of R and K, allowing for 2 missed cleavages, peptide length of 7-50, and a mass range of 500 to 5,000 Da without database splitting. Methionine oxidation was set as a variable modification with a maximum of 3 variable mods per peptide. To include Bpa-containing peptides in the search, a +88.03 Da variable modification was allowed on tyrosine. Cysteine carbamidomethylation was set as a fixed modification. Peptide hits were filtered with a 1% FDR. LFQ was performed by IonQuant, with matching and normalizing between runs.

For both log-phase growth experiments and tunicamycin-treated experiments, FragPipe results, exported as CSV files, were analyzed using R. Significance between conditions was calculated using a two-sided paired t-test and protein intensity ratios were calculated using averaged intensities across four biological replicates. Base-2 fold-change ratios and significance for all proteins and subsets of protein hits were plotted using R.

### Monitoring tunicamycin-induced degradation

The Yeast TAP-tagged ORF collection (Horizon Discovery ^50^) was used to monitor the tunicamycin-induced degradation of protein substrates from our tunicamycin dataset. Strains with the relevant TAP-tagged genes were streaked out on YPD plates and a single colony was used to create a starter culture. The starter culture was diluted to a OD_600_ of 0.1 in a 50 mL YPD culture, and cells were grown at 30 °C until reaching a OD_600_ of ∼ 0.8. The culture was then split into four 5 mL culture tubes. Bortezomib (50 μM final) and tunicamycin (2 ug/mL final) were added from stock solutions to the tubes, and cells were grown for 2 hours. Cells were harvested, washed, and lysed with glass bead beating. Lysates were run on SDS-PAGE and transferred to a PVDF membrane for Western blotting. The membranes were blotted with a single HRP-fused antibody (Santa Cruz Biotechnology, sc-25778) that binds the loading-control protein (GAPDH) and the protein A domain of the TAP-tagged protein.

### RNA extraction and RT-PCR

To verify the induction of ER stress upon addition of tunicamycin, Hac1 mRNA splicing was analyzed by growing cells as in crosslinking experiment, adding 2 μg/mL tunicamycin or 5 mM DTT, incubating for 2 hours, and harvesting approximately 15 OD units of cells. After bead-lysis, RNA was extracted using the PureLink RNA Mini Kit (Thermo). Total RNA was visualized via formaldehyde gel electrophoresis to check for extraction. Extracted RNA was simultaneously converted into DNA and PCR amplified to check for Hac1 splicing using the SuperScript IV UniPrime One-Step RT-PCR system (Thermo) with two sets of primers Hac1_FWD/REV1 and Hac1_FWD/REV2 (See Primers Table below). RT-PCR products were run on an 0.8% agarose gel that was then stained with SYBR Safe (Thermo) and visualized in a ChemiDoc MP (Bio-Rad).

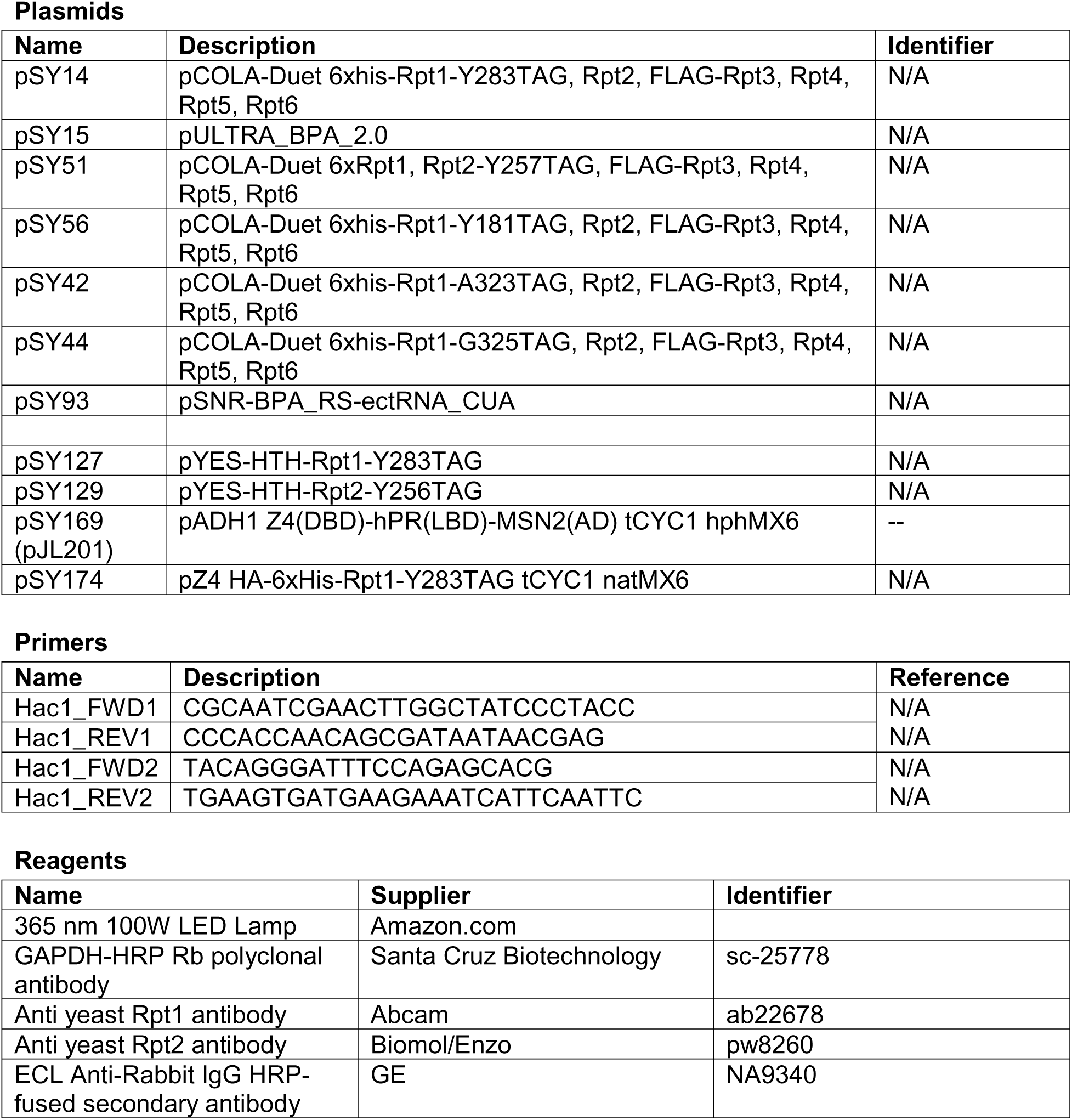

## Notes

### Competing Interest Statement

The authors have declared no competing interest.

### Summary of Updates

The revision added the missing supplement

