## Supplementary Figures for "Identifying endogenous substrates of the 26S proteasome through site-specific photocrosslinking"

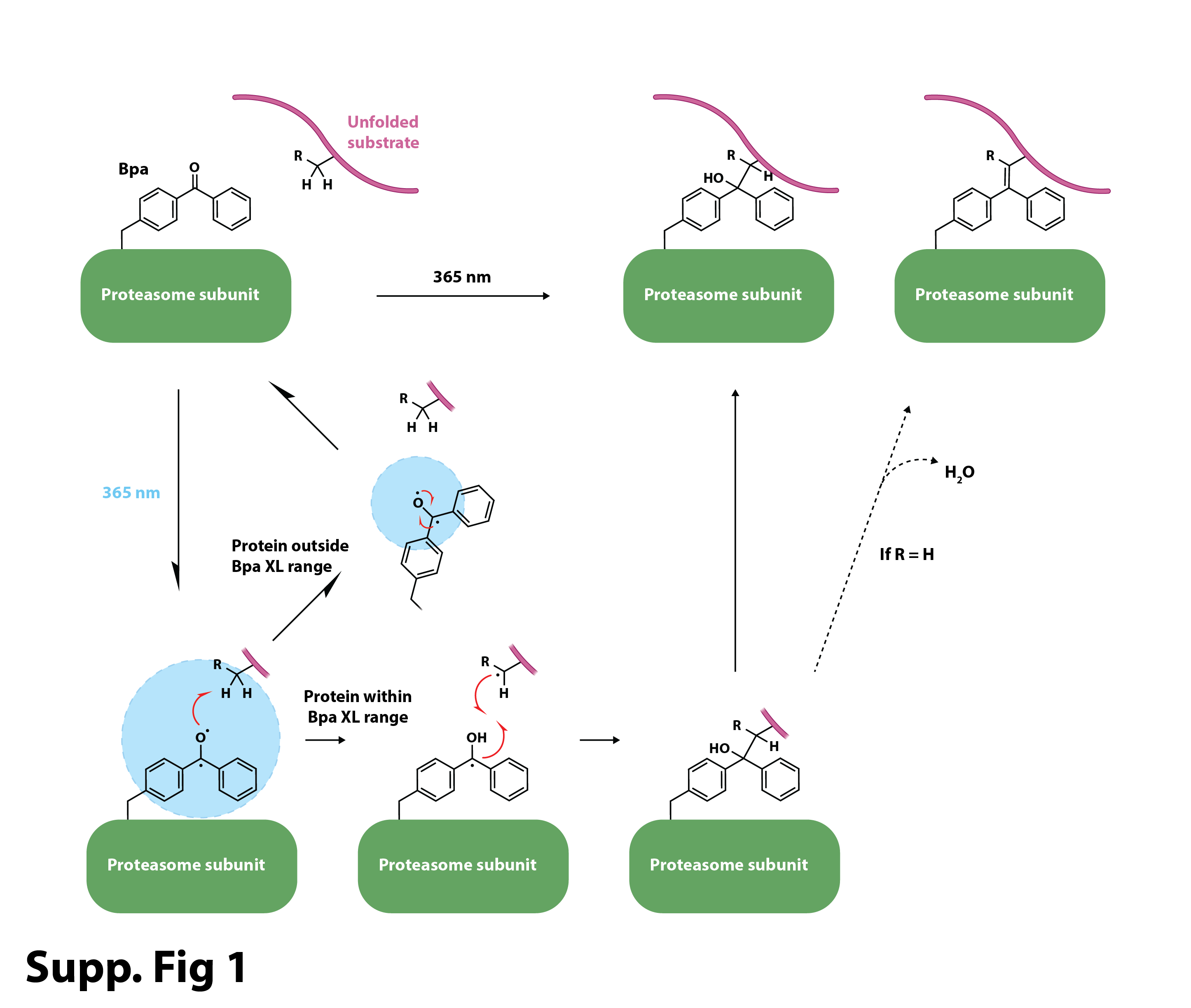

**Supplementary Figure 1. Schematic for the photo-induced crosslinking of 4-benzoyl-L-phenylalaine (Bpa)-containing proteasomes to substrates.** When there is no protein within the crosslinking (XL) range, photoactivated Bpa decays and can be reactivated (cycle on the left).

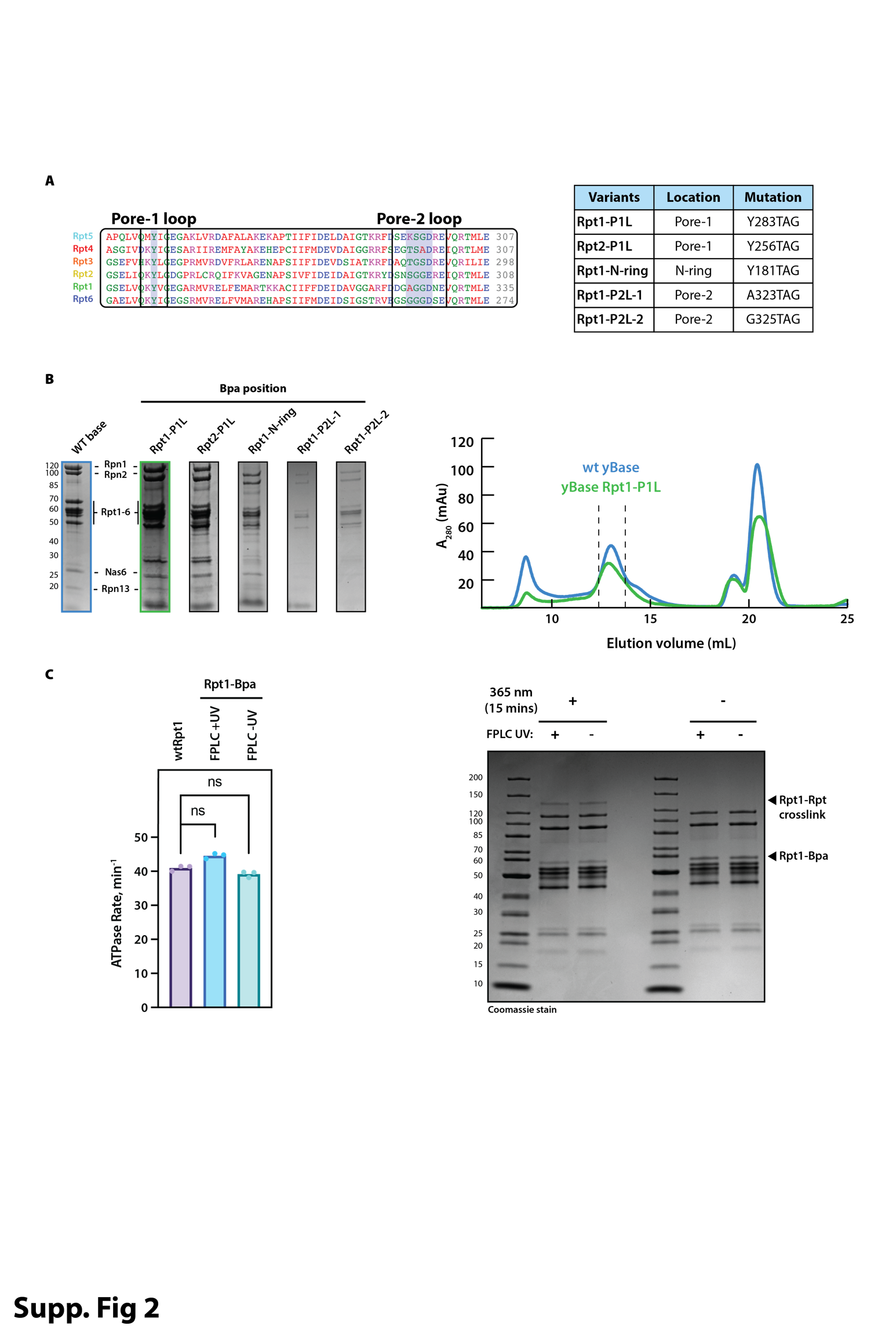

**Supplementary Figure 2. Bpa-containing variants of the proteasome base subcomplex and their basic *in vitro* characterization. A.** Right: Sequence alignment of the Rpt ATPase subunits of the yeast 26S proteasome, with the pore-1 and pore-2 loop motifs boxed and critical residues in these loops shaded in blue. Left: Base variants with amber stop codons in pore-1 or pore-2 loops that were purified, assembled into holoenzymes, and tested. **B.** Left: Coomassie-stained SDS-PAGE gels of variants listed in panel A. Right: Representative size-exclusion chromatography traces for the purification of wild-type and Rpt1-P1L base subcomplexes. **C.** The integrity and activity of Bpa-containing base variants are not affected by the UV-detection during FPLC purification. Left: ATPase rates of wild-type and Rpt1-P1L-Bpa containing base subcomplexes. Right: Rpt1-P1L-Bpa containing base only crosslinks when exposed to 365 nm light and forms an Rpt1-Rpt crosslinking product in the absence of substrate.

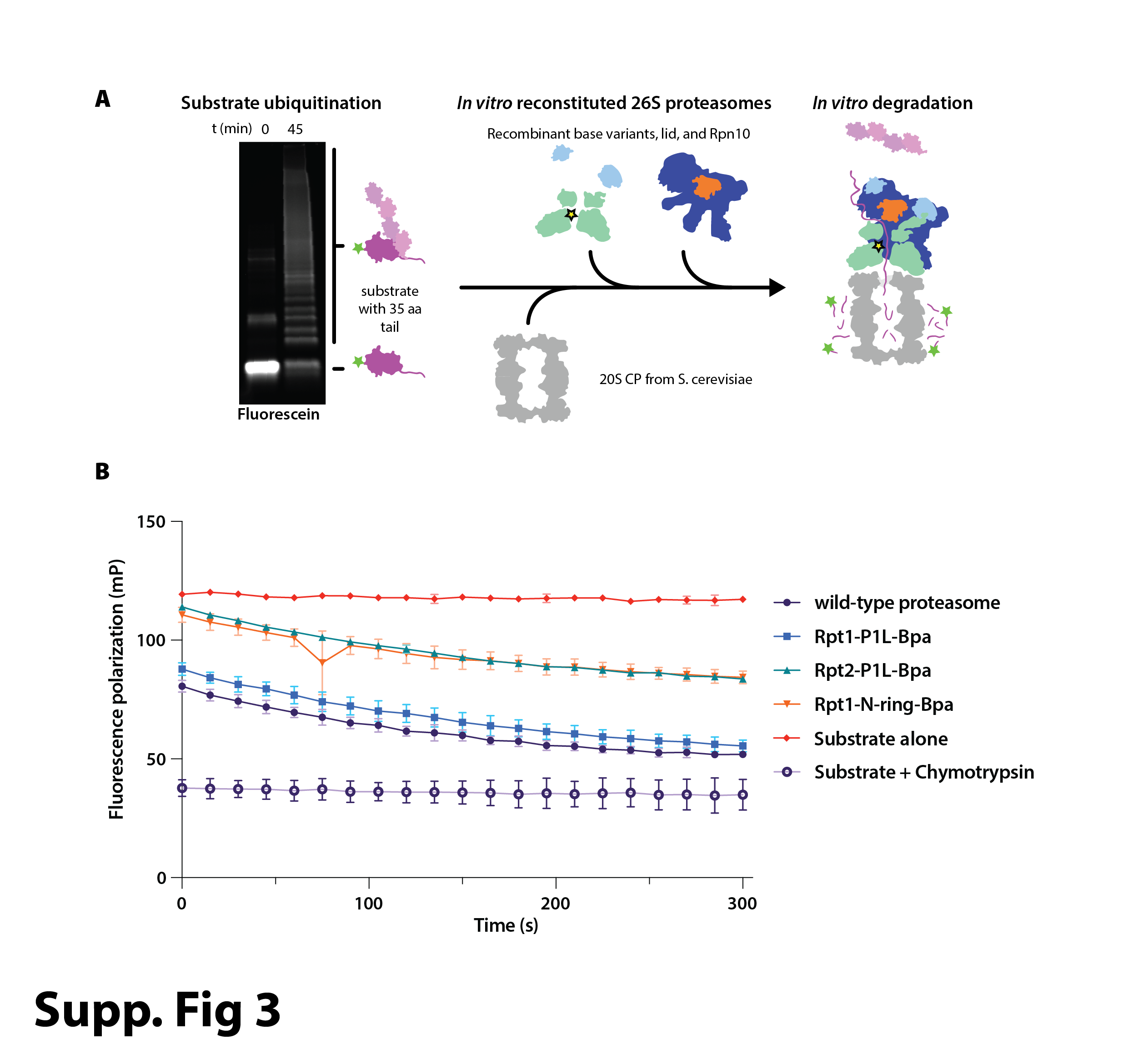

**Supplementary Figure 3. Substrate-degradation assays with reconstituted Bpa-containing 26S holoenzyme. A.** Schematic of *in vitro* reconstitution of 26S proteasome-mediated degradation with Bpa-containing motor subcomplexes. Left: Fluorescence-imaged SDS-PAGE gel for the poly-ubiquitination of the fluorescein-labeled titin model substrate containing a 35-residue flexible tail. Middle: 26S proteasomes are reconstituted from recombinant base, lid, and Rpn10, as well as 20S CP purified from yeast. Right: *In vitro* degradation of the model substrate leads to the release of small fluorescein-labeled peptides and a corresponding decrease in fluorescence polarization. **B.** Averaged traces for the decrease in fluorescence polarization during substrate degradation by wild-type and Bpa-containing proteasome variants. Substrate alone and completely cleaved substrate in the presence of chymotrypsin are shown as controls.

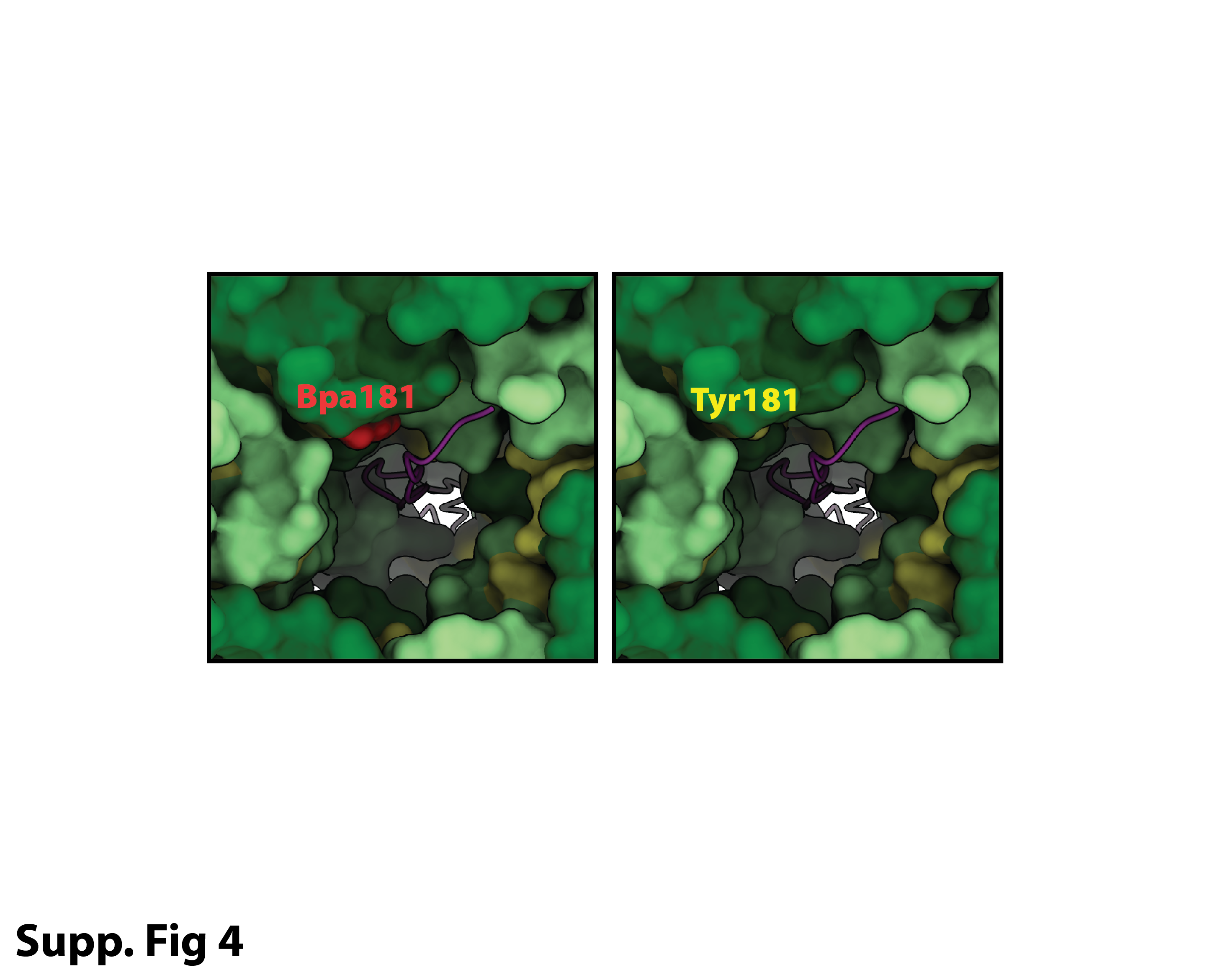

**Supplementary Figure 4. Model of Bpa in the Rpt1 N-ring position.** The ATPase hexamer of the yeast 26S proteasome (PDB ID: 6EF3) is shown in surface representation from the top of the hexameric ring, with the polypeptide backbone of an engaged substrate shown in purple. Tyrosine residues are shown in yellow, and the model for a Tyr181-to-Bpa substitution in the N-terminal domain of Rpt1 is shown in red on the left.

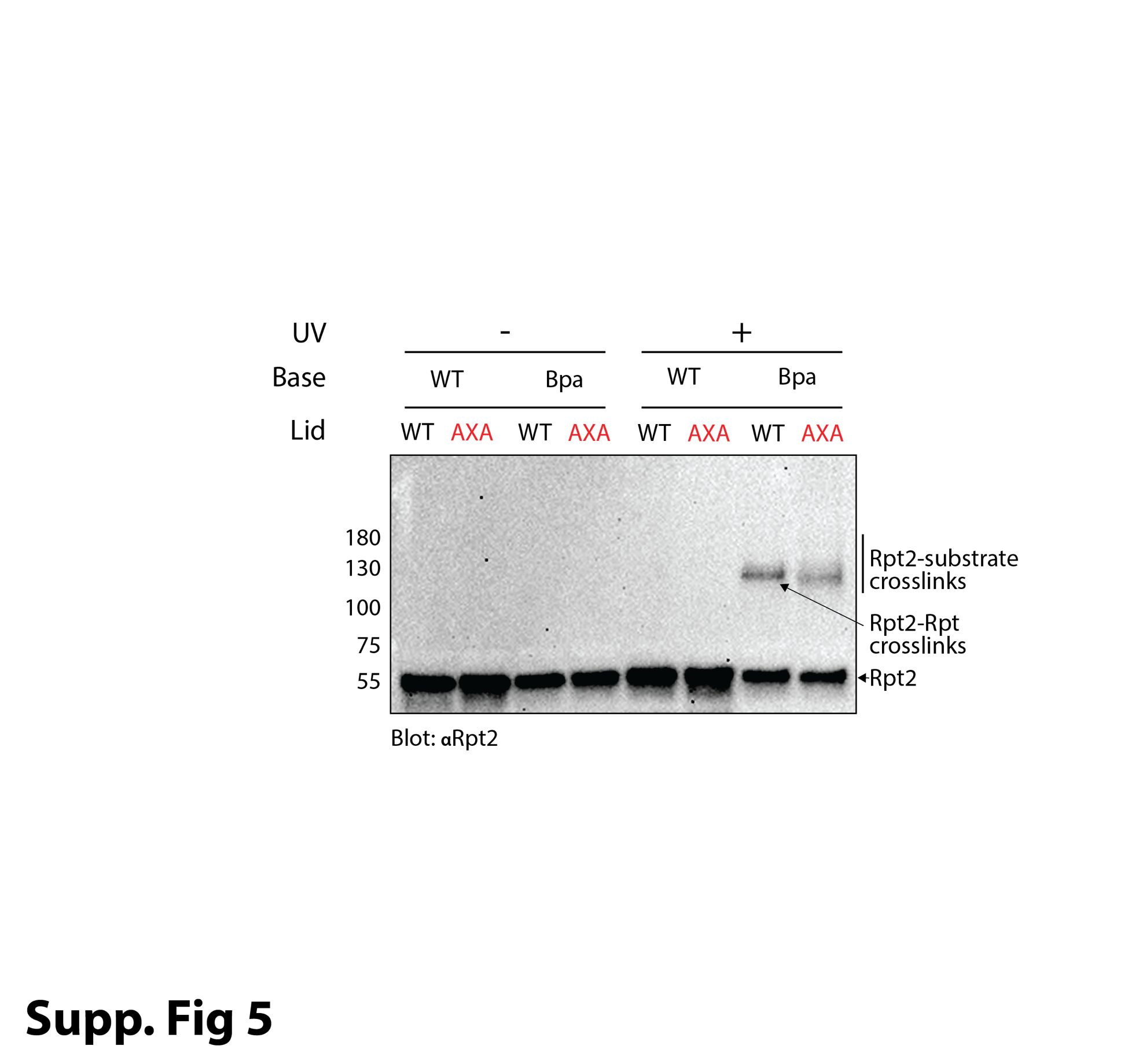

**Supplementary Figure 5. Crosslinking of reconstituted proteasomes containing Rpt2-PL1-Bpa.** Shown is the anti-Rpt2 Western blot of samples from titin-substrate degradation reactions performed with 26S proteasomes that were reconstituted with either wild-type or Rpt2-PL1-Bpa-containing base subcomplexes and wild-type or Rpn11-AXA mutant lid subcomplexes, in the absence or presence of UV light illumination. Stalling of the ubiquitinated substrate with Rpn11-AXA leads to a Rpt2-substrate crosslink, whereas uninhibited deubiquitination and thus rapid degradation increases the extent of Rpt2-Rpt intra-base crosslinks.

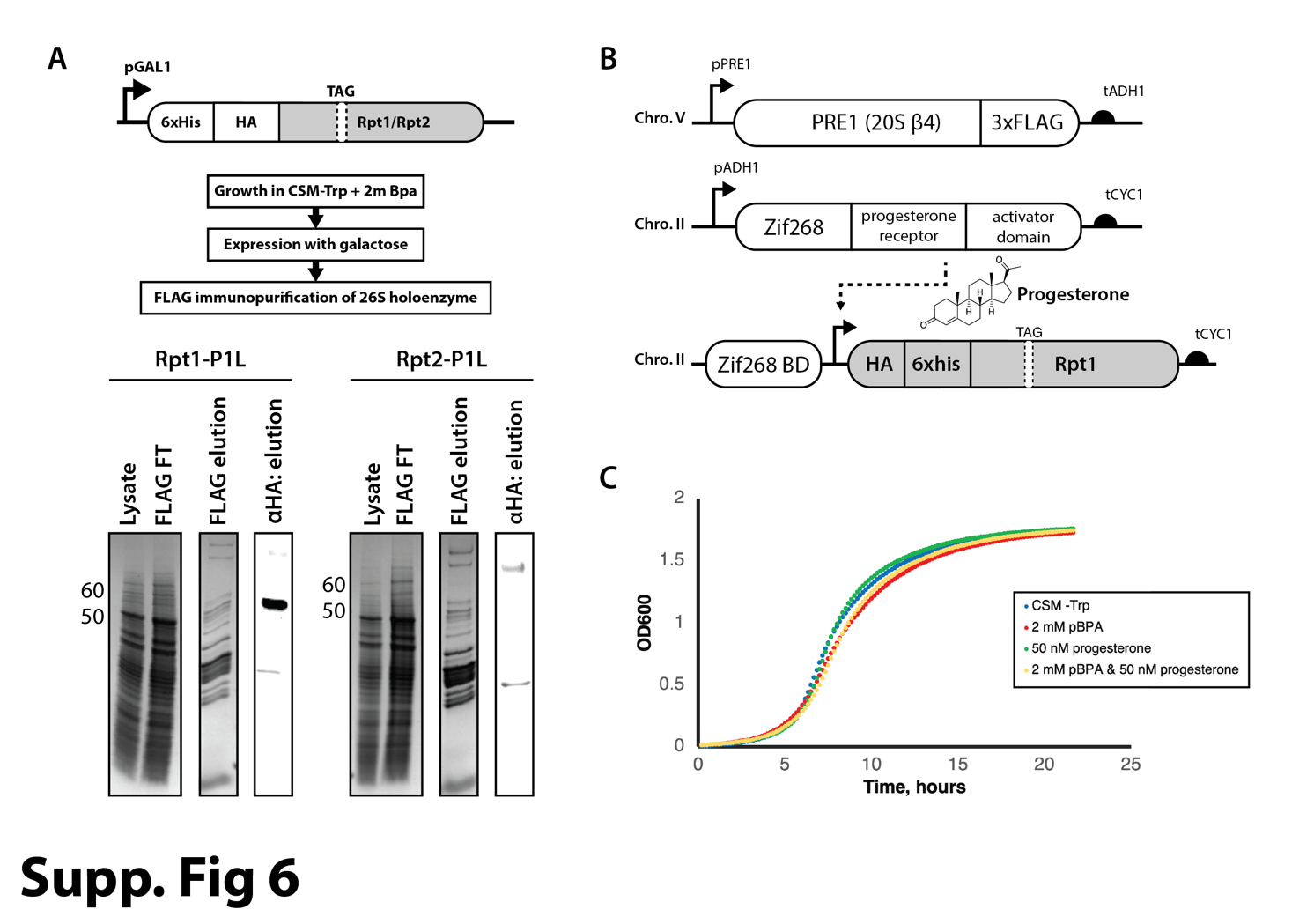

**Supplementary Figure 6. Expression and proteasome incorporation of Rpt1-PL1-Bpa and Rpt2-PL1-Bpa in yeast cells. A.** Top: Schematic for the galactose-inducible expression of Bpa-containing Rpt1 and Rpt2. Bottom: Coomassie-stained SDS-PAGE gels for the yeast lysate, FLAG flowthrough, and FLAG elution of Rpt1-PL1-Bpa (lanes 1-3) and Rpt2-PL1-Bpa lanes 5-7, as well as the corresponding anti-HA Western blot of the FLAG elution samples (lane 4, 8) show that Rpt2-Bpa is not efficiently incorporated into 26S proteasomes in live cells. **B.** Schematic of the orthogonal, progesterone-dependent expression system. **C.** Growth curves show no defect upon the addition of Bpa (pBPA), the expression of Rpt1-PL1-Bpa after addition of progesterone, or both.

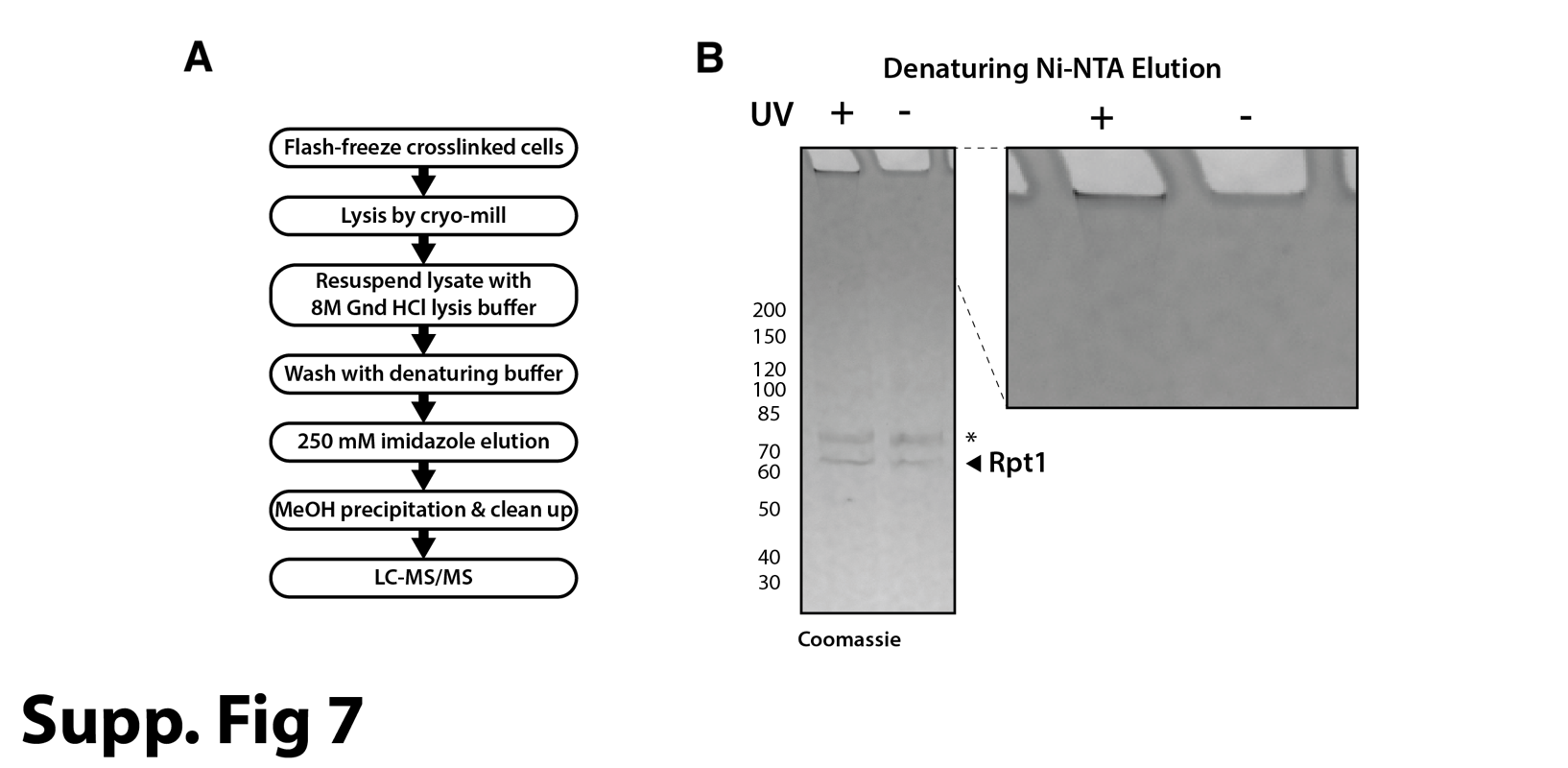

**Supplementary Figure 7. Expression of Bpa-containing Rpt1 in yeast cells. A.** Scheme of experimental procedures for denaturing Ni-NTA purification and LC-MS/MS analysis of His-tagged Rpt1-PL1-Bpa and crosslinked substrates. **B.** Coomassie-stained SDS PAGE gel of elution samples for denaturing Ni-NTA purification of Rpt1-PL1-Bpa from cells with or without UV exposure.

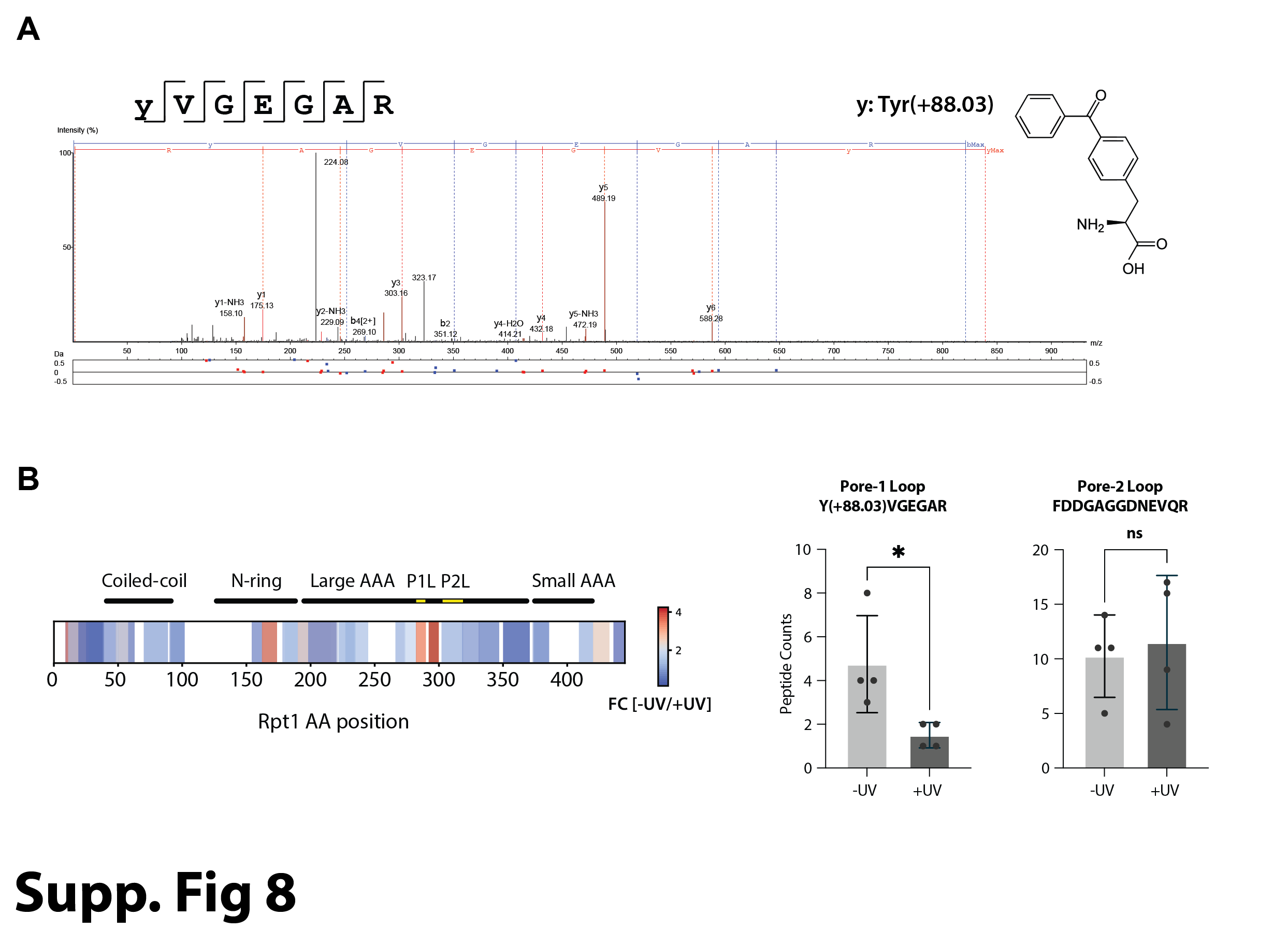

**Supplementary Figure 8. Mass-spectrometric analysis of Bpa-containing Rpt1. A.** Mass spectrum of Rpt1’s Bpa-containing pore-1 loop peptide. The y in the yVGEGAR sequence corresponds to a tyrosine residue with a mass shift of +88.03 Daltons, which is the mass difference between Tyr and Bpa. **B.** Left: Analysis of fold enrichment for the intensities of all Rpt1-derived peptides, indicating that peptides encompassing Rpt1’s pore-1 loop are depleted in the +UV conditions. Colors represent average fold enrichment of peptides in -UV over +UV-exposed cells for n = 4 biological replicates. Right: Quantification of pore-1-loop and pore-2-loop-derived peptides by spectral counts reveals a depletion of the pore-1 loop peptide for the sample purified from UV-exposed cells. Shown are the mean values and standard deviations of the mean for n = 4 biological replicates. Statistical significance was calculated using a one-way ANOVA test: * p < 0.05; ns, p > 0.05.

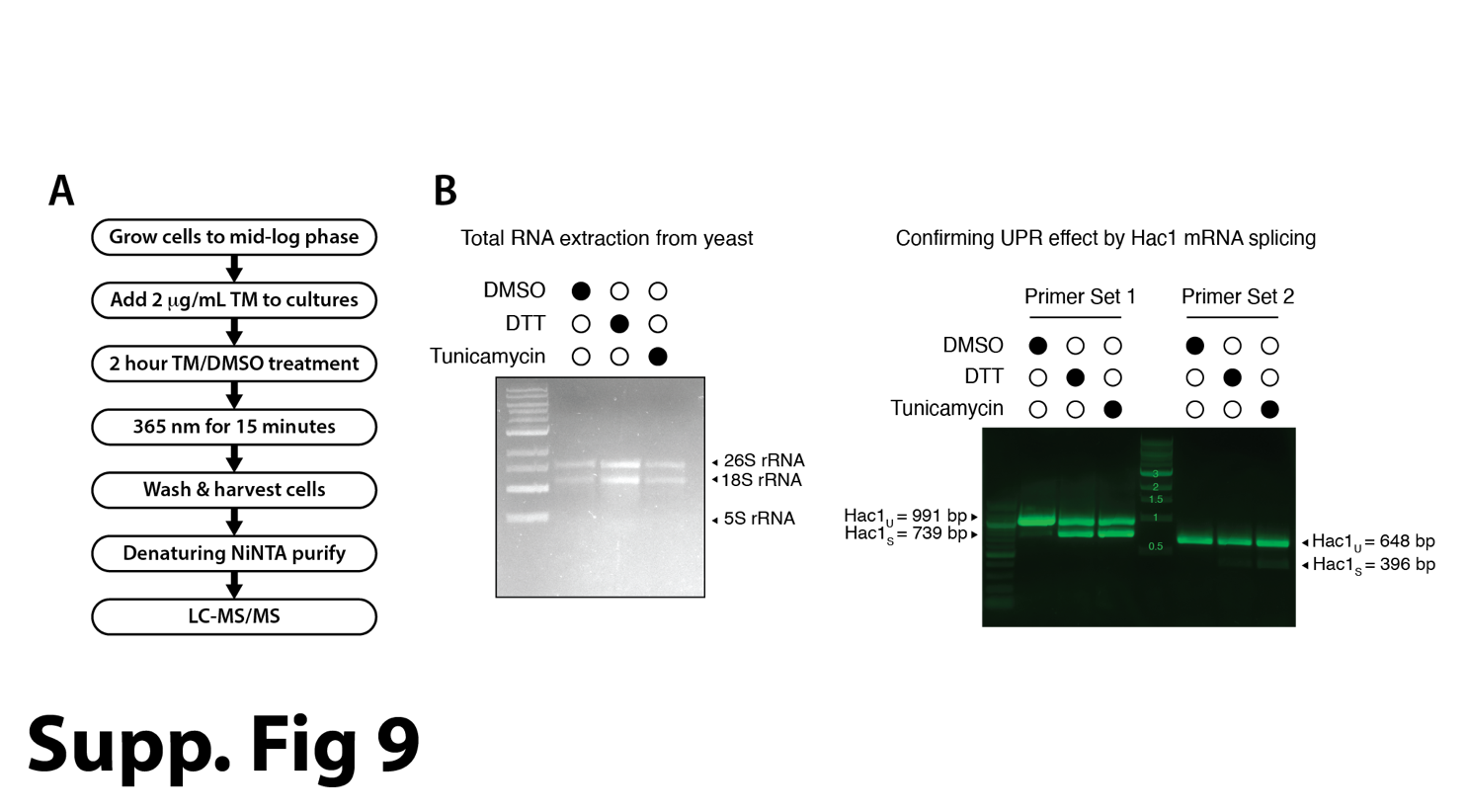

**Supplementary Figure 9. Confirmation of the tunicamycin-inducted unfolded protein response (UPR) by RT-PCR. A.** Scheme of experimental procedures for the proteasome crosslinking and purification of tunicamycin-induced substrates. **B.** To confirm the effects of tunicamycin additions, RNA was extracted from cells (left) and splicing of the mRNA for Hac1 was assessed by RT-PCR with two sets of primers (right). Addition of DTT to cells was used as a positive control for Hac1 splicing.

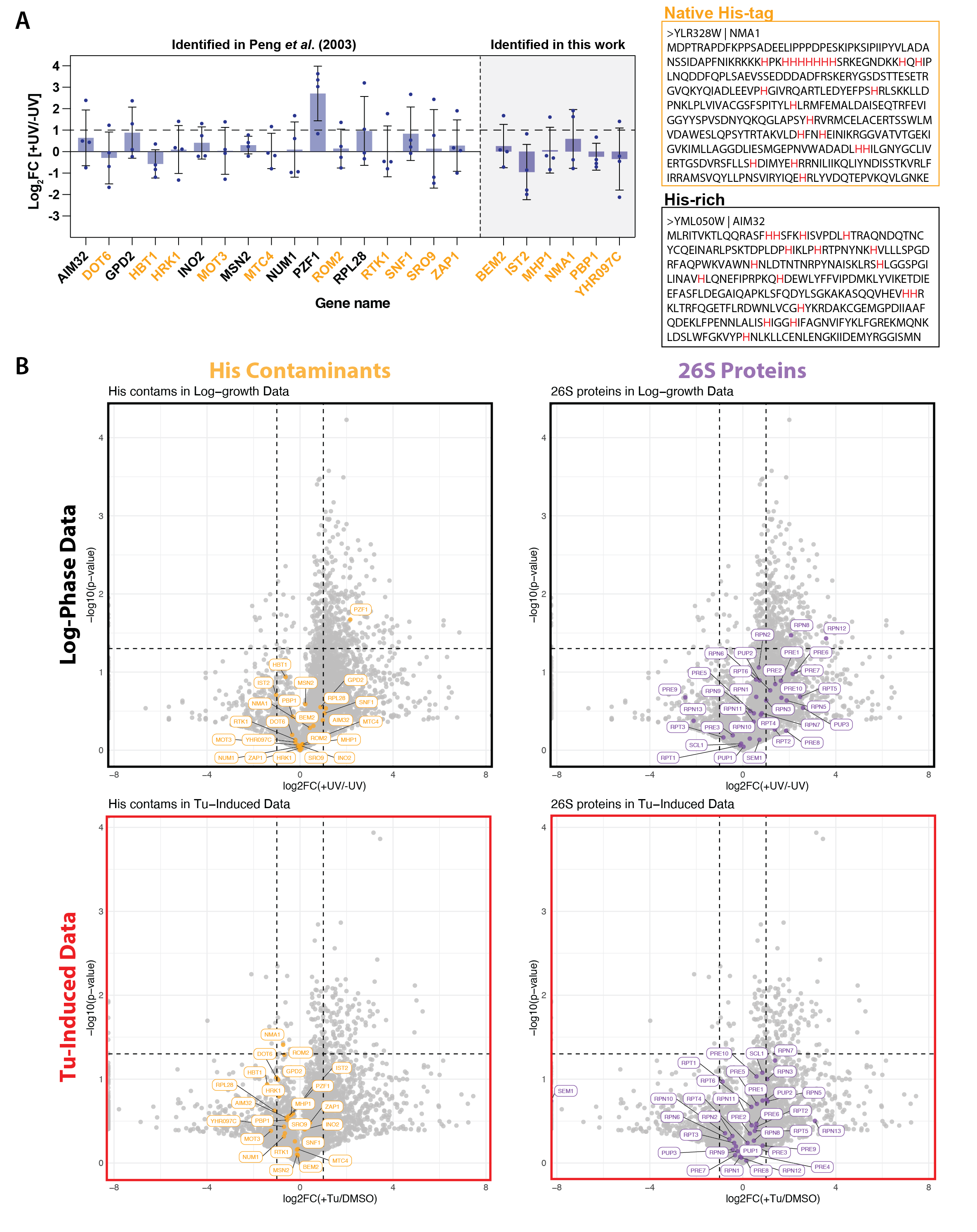

**Supplementary Figure 10. Comparison of abundances for His-containing contaminants and 26S proteasome subunits in mass-spec data for cells in log-phase growth and under tunicamycin-induced stress. A.** Left: Log_2_-fold changes (FC) of histidine-rich proteins that co-purify under denaturing conditions with His_6_ -Rpt1-Bpa from the log-phase growth experiment. These proteins include 17 previously identified contaminants of Ni-NTA affinity purifications ^27^ and six newly identified proteins. Right: Representative sequences for a natively His-tagged protein, NMA1, and a His-rich protein, AIM32. **B.** Volcano plots for the changes in protein abundances derived from mass-spec analyses of samples from log-phase growth and Tu-stressed cells. For each dataset His-contaminants are labelled in yellow and 26S proteasome subunits are labelled in purple.
